# Meta-analysis of genetic association with diagnosed Alzheimer’s disease identifies novel risk loci and implicates Abeta, Tau, immunity and lipid processing

**DOI:** 10.1101/294629

**Authors:** BW Kunkle, B Grenier-Boley, R Sims, JC Bis, AC Naj, A Boland, M Vronskaya, SJ van der Lee, A Amlie-Wolf, C Bellenguez, A Frizatti, V Chouraki, ER Martin, K Sleegers, N Badarinarayan, J Jakobsdottir, KL Hamilton-Nelson, R Aloso, R Raybould, Y Chen, AB Kuzma, M Hiltunen, T Morgan, S Ahmad, BN Vardarajan, J Epelbaum, P Hoffmann, M Boada, GW Beecham, JG Garnier, D Harold, AL Fitzpatrick, O Valladares, ML Moutet, A Gerrish, AV Smith, L Qu, D Bacq, N Denning, X Jian, Y Zhao, MD Zompo, NC Fox, ML Grove, SH Choi, I Mateo, JT Hughes, HH Adams, J Malamon, FS Garcia, Y Patel, JA Brody, B Dombroski, MCD Naranjo, M Daniilidou, G Eiriksdottir, S Mukherjee, D Wallon, J Uphill, T Aspelund, LB Cantwell, F Garzia, D Galimberti, E Hofer, M Butkiewics, B Fin, E Scarpini, C Sarnowski, W Bush, S Meslage, J Kornhuber, CC White, Y Song, RC Barber, S Engelborghs, S Pichler, D Voijnovic, PM Adams, R Vandenberghe, M Mayhaus, LA Cupples, MS Albert, PP De Deyn, W Gu, JJ Himali, D Beekly, A Squassina, AM Hartmann, A Orellana, D Blacker, E Rodriguez-Rodriguez, S Lovestone, ME Garcia, RS Doody, CM Fernadez, R Sussams, H Lin, TJ Fairchild, YA Benito, C Holmes, H Comic, MP Frosch, H Thonberg, W Maier, G Roschupkin, B Ghetti, V Giedraitis, A Kawalia, S Li, RM Huebinger, L Kilander, S Moebus, I Hernández, MI Kamboh, R Brundin, J Turton, Q Yang, MJ Katz, L Concari, J Lord, AS Beiser, CD Keene, S Helisalmi, I Kloszewska, WA Kukull, AM Koivisto, A Lynch, L Tarraga, EB Larson, A Haapasalo, B Lawlor, TH Mosley, RB Lipton, V Solfrizzi, M Gill, WT Longstreth, TJ Montine, V Frisardi, S Ortega-Cubero, F Rivadeneira, RC Petersen, V Deramecourt, A Ciaramella, E Boerwinkle, EM Reiman, N Fievet, C Caltagirone, JI Rotter, JS Reisch, O Hanon, C Cupidi, AG Uitterlinden, DR Royall, C Dufouil, RG Maletta, S Moreno-Grau, M Sano, A Brice, R Cecchetti, P St George-Hyslop, K Ritchie, M Tsolaki, DW Tsuang, B Dubois, D Craig, CK Wu, H Soininen, D Avramidou, RL Albin, L Fratiglioni, A Germanou, LG Apostolova, L Keller, M Koutroumani, SE Arnold, F Panza, O Gkatzima, S Asthana, D Hannequin, P Whitehead, CS Atwood, P Caffarra, H Hampel, CT Baldwin, L Lannfelt, DC Rubinsztein, LL Barnes, F Pasquier, L Frölich, S Barral, B McGuinness, TG Beach, JI Johnston, JT Becker, P Passmore, EH Bigio, JM Schott, TD Bird, JD Warren, BF Boeve, MK Lupton, JD Bowen, P Proitsi, A Boxer, JF Powell, JR Burke, JK Kauwe, JM Burns, M Mancuso, JD Buxbaum, U Bonuccelli, NJ Cairns, A McQuillin, C Cao, G Livingston, CS Carlson, NJ Bass, CM Carlsson, J Hardy, RM Carney, J Bras, MM Carrasquillo, R Guerreiro, M Allen, HC Chui, E Fisher, DH Cribbs, C Masullo, EA Crocco, C DeCarli, G Bisceglio, M Dick, L Ma, R Duara, NR Graff-Radford, DA Evans, A Hodges, KM Faber, M Scherer, KB Fallon, M Riemenschneider, DW Fardo, R Heun, MR Farlow, S Ferris, M Leber, TM Foroud, I Heuser, DR Galasko, I Giegling, M Gearing, M Hüll, DH Geschwind, JR Gilbert, J Morris, RC Green, K Mayo, JH Growdon, T Feulner, RL Hamilton, LE Harrell, D Drichel, LS Honig, TD Cushion, MJ Huentelman, P Hollingworth, CM Hulette, BT Hyman, R Marshall, GP Jarvik, A Meggy, E Abner, G Menzies, LW Jin, G Leonenko, G Jun, D Grozeva, A Karydas, G Russo, JA Kaye, R Kim, F Jessen, NW Kowall, B Vellas, JH Kramer, E Vardy, FM LaFerla, KH Jöckel, JJ Lah, M Dichgans, JB Leverenz, D Mann, AI Levey, S Pickering-Brown, AP Lieberman, N Klopp, KL Lunetta, HE Wichmann, CG Lyketsos, K Morgan, DC Marson, K Brown, F Martiniuk, C Medway, DC Mash, MM Nöthen, E Masliah, NM Hooper, WC McCormick, A Daniele, SM McCurry, A Bayer, AN McDavid, J Gallacher, AC McKee, H van den Bussche, M Mesulam, C Brayne, BL Miller, S Riedel-Heller, CA Miller, JW Miller, A Al-Chalabi, JC Morris, CE Shaw, AJ Myers, J Wiltfang, S O’Bryant, E Coto, JM Olichney, V Alvarez, JE Parisi, AB Singleton, HL Paulson, J Collinge, W Perry, S Mead, E Peskind, M Rosser, A Pierce, N Ryan, WW Poon, B Nacmias, H Potter, S Sorbi, JF Quinn, E Sacchinelli, A Raj, G Spalletta, M Raskind, P Bossù, B Reisberg, R Clarke, C Reitz, AD Smith, JM Ringman, D Warden, ED Roberson, G Wilcock, E Rogaeva, AC Bruni, HJ Rosen, M Gallo, RN Rosenberg, Y Ben-Shlomo, MA Sager, P Mecocci, AJ Saykin, P Pastor, ML Cuccaro, JM Vance, JA Schneider, LS Schneider, WW Seeley, AG Smith, JA Sonnen, S Spina, RA Stern, RH Swerdlow, RE Tanzi, JQ Trojanowski, JC Troncoso, VM Van Deerlin, LJ Van Eldik, HV Vinters, JP Vonsattel, S Weintraub, KA Welsh-Bohmer, KC Wilhelmsen, J Williamson, TS Wingo, RL Woltjer, CB Wright, CE Yu, L Yu, Alzheimer Disease Genetics Consortium (ADGC), The European Alzheimer’s Disease Initiative (EADI), Cohorts for Heart and Aging Research in Genomic Epidemiology Consortium (CHARGE), Genetic and Environmental Risk in AD/Defining Genetic, Polygenic and Environmental Risk for Alzheimer’s Disease Consortium (GERAD/PERADES), PK Crane, DA Bennett, V Boccardi, PL De Jager, N Warner, OL Lopez, S McDonough, M Ingelsson, P Deloukas, C Cruchaga, C Graff, R Gwilliam, M Fornage, AM Goate, P Sanchez-Juan, PG Kehoe, N Amin, N Ertekin-Taner, C Berr, S Debette, S Love, LJ Launer, SG Younkin, JF Dartigues, C Corcoran, MA Ikram, DW Dickson, D Campion, J Tschanz, H Schmidt, H Hakonarson, R Munger, R Schmidt, LA Farrer, C Van Broeckhoven, MC O’Donovan, AL DeStefano, L Jones, JL Haines, JF Deleuze, MJ Owen, V Gudnason, R Mayeux, V Escott-Price, BM Psaty, A Ruiz, A Ramirez, LS Wang, CM van Duijn, PA Holmans, S Seshadri, J Williams, P Amouyel, GD Schellenberg, JC Lambert, MA Pericak-Vance

**Author notes:** equal contribution first author (BW Kunkle, B Grenier-Boley). equal contribution senior author (CM van Duijn, PA Holmans, S Seshadri, J Williams, P Amouyel, GD Schellenberg, JC Lambert, MA Pericak-Vance). corresponding author Materials and Correspondence: Brian W. Kunkle, PhD, MPH Hussman Institute for Human Genomics Miller School of Medicine University of Miami 1501 NW 10^th^ Ave Miami, FL 33136 Jean-Charles Lambert, PhD INSERM, U1167 Laboratoire d’Excellence Distalz Institut Pasteur de Lille University Lille F59000 Lille, France Margaret Pericak-Vance, PhD Hussman Institute for Human Genomics Miller School of Medicine University of Miami 1501 NW 10^th^ Ave Miami, FL 33136.

## Abstract

Late-onset Alzheimer’s disease (LOAD, onset age > 60 years) is the most prevalent dementia in the elderly^1^, and risk is partially driven by genetics^2^. Many of the loci responsible for this genetic risk were identified by genome-wide association studies (GWAS)^3–8^. To identify additional LOAD risk loci, the we performed the largest GWAS to date (89,769 individuals), analyzing both common and rare variants. We confirm 20 previous LOAD risk loci and identify four new genome-wide loci (*IQCK*, *ACE*, *ADAM10*, and *ADAMTS1*). Pathway analysis of these data implicates the immune system and lipid metabolism, and for the first time tau binding proteins and APP metabolism. These findings show that genetic variants affecting APP and Aβ processing are not only associated with early-onset autosomal dominant AD but also with LOAD. Analysis of AD risk genes and pathways show enrichment for rare variants (*P* = 1.32 × 10^−7^) indicating that additional rare variants remain to be identified.

## Main Text

Our previous work identified 19 genome-wide significant common variant signals in addition to *APOE*^9^, that influence risk for LOAD. These signals, combined with ‘subthreshold’ common variant associations, account for ~31% of the genetic variance of LOAD^2^, leaving the majority of genetic risk uncharacterized^10^. To search for additional signals, we conducted a GWAS meta-analysis of non-Hispanic Whites (NHW) using a larger sample (17 new, 46 total datasets) from our group, the International Genomics of Alzheimer’s Project (IGAP) (composed of four AD consortia: ADGC, CHARGE, EADI, and GERAD). This sample increases our previous discovery sample (Stage 1) by 29% for cases and 13% for controls (N=21,982 cases; 41,944 controls) (**Supplementary Table 1** and **2**, and **Supplementary Note**). To sample both common and rare variants (minor allele frequency MAF ≥ 0.01, and MAF < 0.01, respectively), we imputed the discovery datasets using a 1000 Genomes reference panel consisting of 36,648,992 single-nucleotide variants, 1,380,736 insertions/deletions, and 13,805 structural variants. After quality control, 9,456,058 common variants and 2,024,574 rare variants were selected for analysis (a 63% increase from our previous common variant analysis in 2013). Genotype dosages were analyzed within each dataset, and then combined with meta-analysis (**Supplementary Figures 1 and 2** and **Supplementary Table 3**). The Stage 1 discovery meta-analysis was first followed by Stage 2 using the I-select chip we previously developed in Lambert et al (including 11,632 variants, N=18,845) and finally stage 3A (N=6,998). The final sample was 33,692 clinical AD cases and 56,077 controls.

Meta-analysis of Stages 1 and 2 produced 21 associations with *P ≤* 5×10^−8^ (**Table 1** and **Figure 1**). Of these, 18 were previously reported as genome-wide significant and three of them are signals not initially described in Lambert et al: the rare R47H *TREM2* coding variant previously reported by others^11–13^; *ECDH3* (rs7920721) which was recently identified as a potential genome-wide significant AD risk locus in several studies^23–25^ and *ACE* (rs138190086). In addition, four signal showed suggestive association with a P-value<5.10^−7^ (respectively rs593742, rs830500, rsrs7295246 and rs7185636 for *ADAM10, ADAMTS1, ADAMTS20, and IQCK*).

**Table 1.** Summary of discovery stage 1, stage 2 and overall meta-analyses results for identified loci reaching genome-wide significance after stages 1 and 2.

|  |  |  |  |  |  | Stage 1 Discovery (n=63,926) |  | Stage 2 (n=18,845) |  | Overall Stages 1 + Stage 2 (n=82,771) |  |  |
| --- | --- | --- | --- | --- | --- | --- | --- | --- | --- | --- | --- | --- |
| SNP<sup>a</sup> | Chr. | Position<sup>b</sup> | Closest gene<sup>c</sup> | Major/<br>minor alleles | MAF<sup>d</sup> | OR (95% CI)<sup>e</sup> | Meta P<br>value | OR (95% CI)<sup>e</sup> | Meta P value | OR (95% CI)<sup>e</sup> | Meta P value | I<sup>2</sup> (%),<br>P value<sup>f</sup> |
| Previous genome-wide significant loci still reaching significance |  |  |  |  |  |  |  |  |  |  |  |  |
| rs4844610 | 1 | 207802552 | CR1 | C/A | 0.187 | 1.16 (1.12-1.20) | 8.2 x 10<sup>-16</sup> | 1.20 (1.13-1.27) | 3.8 x 10<sup>-10</sup> | 1.17 (1.13-1.21) | 3.6 x 10<sup>-24</sup> | 0, 8 x 10<sup>-1</sup> |
| rs6733839 | 2 | 127892810 | BIN1 | C/T | 0.407 | 1.18 (1.15-1.22) | 4.0 x 10<sup>-28</sup> | 1.23 (1.18-1.29) | 2.0 x 10<sup>-18</sup> | 1.20 (1.17-1.23) | 2.1 x 10<sup>-44</sup> | 15, 2 x 10<sup>-1</sup> |
| rs10933431 | 2 | 233981912 | INPP5D | C/G | 0.223 | 0.90 (0.87-0.94) | 2.6 x 10<sup>-7</sup> | 0.92 (0.87-0.97) | 3.2 x 10<sup>-3</sup> | 0.91 (0.88-0.94) | 3.4 x 10<sup>-9</sup> | 0, 8 x 10<sup>-1</sup> |
| rs78738018 | 6 | 32575406 | HLA-DQB1 | T/A | 0.270 | 1.10 (1.06-1.14) | 5.1 x 10<sup>-8</sup> | 1.11 (1.06-1.17) | 5.7 x 10<sup>-5</sup> | 1.10 (1.07-1.13) | 1.4 x 10<sup>-11</sup> | 10, 3 x 10<sup>-1</sup> |
| rs75932628 | 6 | 41129252 | TREM2 | C/T | 0.008 | 2.01 (1.65-2.44) | 2.9 x 10<sup>-12</sup> | 2.50 (1.56-4.00) | 1.5 x 10<sup>-4</sup> | 2.08 (1.73-2.49) | 2.7 x 10<sup>-15</sup> | 0, 6 x 10<sup>-1</sup> |
| rs9473117 | 6 | 47431284 | CD2AP | A/C | 0.280 | 1.09 (1.05-1.12) | 2.3 x 10<sup>-7</sup> | 1.11 (1.05-1.16) | 1.0 x 10<sup>-4</sup> | 1.09 (1.06-1.12) | 1.2 x 10<sup>-10</sup> | 0, 6 x 10<sup>-1</sup> |
| rs12539172 | 7 | 100091795 | NYAP1<sup>g</sup> | C/T | 0.303 | 0.93 (0.91-0.96) | 2.1 x 10<sup>-5</sup> | 0.89 (0.84-0.93) | 2.1 x 10<sup>-6</sup> | 0.92 (0.90-0.95) | 9.3 x 10<sup>-10</sup> | 0, 8 x 10<sup>-1</sup> |
| rs11762262 | 7 | 143107876 | EPHA1 | C/A | 0.199 | 0.90 (0.87-0.94) | 3.1 x 10<sup>-8</sup> | 0.91 (0.86-0.96) | 1.1 x 10<sup>-3</sup> | 0.90 (0.88-0.93) | 1.3 x 10<sup>-10</sup> | 0, 5 x 10<sup>-1</sup> |
| rs73223431 | 8 | 27219987 | PTK2B | C/T | 0.367 | 1.10 (1.07-1.13) | 8.3 x 10<sup>-10</sup> | 1.11 (1.06-1.16) | 1.5 x 10<sup>-5</sup> | 1.10 (1.07-1.13) | 6.3 x 10<sup>-14</sup> | 0, 6 x 10<sup>-1</sup> |
| rs9331896 | 8 | 27467686 | CLU | T/C | 0.387 | 0.88 (0.85-0.91) | 3.6 x 10<sup>-16</sup> | 0.87 (0.83-0.91) | 1.7 x 10<sup>-9</sup> | 0.88 (0.85-0.90) | 4.6 x 10<sup>-24</sup> | 3, 4 x 10<sup>-1</sup> |
| rs3740688 | 11 | 47380340 | SPI1<sup>h</sup> | T/G | 0.448 | 0.91 (0.89-0.94) | 9.7 x 10<sup>-11</sup> | 0.93 (0.88-0.97) | 1.2 x 10<sup>-3</sup> | 0.92 (0.89-0.94) | 5.4 x 10<sup>-13</sup> | 4, 4 x 10<sup>-1</sup> |
| rs7933202 | 11 | 59936926 | MS4A2 | A/C | 0.391 | 0.89 (0.86-0.92) | 2.2 x 10<sup>-15</sup> | 0.90 (0.86-0.95) | 1.6 x 10<sup>-5</sup> | 0.89 (0.87-0.92) | 1.9 x 10<sup>-19</sup> | 27, 5 x 10<sup>-2</sup> |
| rs3851179 | 11 | 85868640 | PICALM | C/T | 0.356 | 0.89 (0.86-0.91) | 5.8 x 10<sup>-16</sup> | 0.85 (0.81-0.89) | 6.1 x 10<sup>-11</sup> | 0.88 (0.86-0.90) | 6.0 x 10<sup>-25</sup> | 0, 8 x 10<sup>-1</sup> |
| rs11218343 | 11 | 121435587 | SORL1 | T/C | 0.040 | 0.81 (0.76-0.88) | 2.7 x 10<sup>-8</sup> | 0.77 (0.68-0.87) | 1.8 x 10<sup>-5</sup> | 0.80 (0.75-0.85) | 2.9 x 10<sup>-12</sup> | 7, 3 x 10<sup>-1</sup> |
| rs17125924 | 14 | 53391680 | FERMT2 | A/G | 0.093 | 1.13 (1.08-1.19) | 6.6 x 10<sup>-7</sup> | 1.15 (1.06-1.25) | 5.0 x 10<sup>-4</sup> | 1.14 (1.09-1.18) | 1.4 x 10<sup>-9</sup> | 8, 3 x 10<sup>-1</sup> |
| rs12881735 | 14 | 92932828 | SLC24A4 | T/C | 0.221 | 0.92 (0.88-0.95) | 4.9 x 10<sup>-7</sup> | 0.92 (0.87-0.97) | 4.3 x 10<sup>-3</sup> | 0.92 (0.89-0.94) | 7.4x 10<sup>-9</sup> | 0, 6 x 10<sup>-1</sup> |
| rs3752246 | 19 | 1056492 | ABCA7 | C/G | 0.182 | 1.13 (1.09-1.18) | 6.6 x 10<sup>-10</sup> | 1.18 (1.11-1.25) | 4.7 x 10<sup>-8</sup> | 1.15 (1.11-1.18) | 3.1 x 10<sup>-16</sup> | 0, 5 x 10<sup>-1</sup> |
| rs429358 | 19 | 45411941 | APOE | T/C | 0.216 | 0.30 (0.28-0.31) | 1.2 x 10<sup>-881</sup> | APOE region not carried forward to replication stage |  |  |  |  |
| rs6024870 | 20 | 54997568 | CASS4 | G/A | 0.088 | 0.88 (0.84-0.93) | 1.1 x 10<sup>-6</sup> | 0.90 (0.82-0.97) | 9.0 x 10<sup>-3</sup> | 0.88 (0.85-0.92) | 3.5 x 10<sup>-8</sup> | 0, 9 x 10<sup>-1</sup> |
| New genome-wide significant loci reaching significance |  |  |  |  |  |  |  |  |  |  |  |  |
| rs138190086 | 7 | 61538148 | ACE | G/A | 0.02 | 1.29 (1.15-1.44) | 7.4 x 10<sup>-6</sup> | 1.41 (1.18-1.69) | 1.8 x 10<sup>-4</sup> | 1.32 (1.20-1.45) | 7.5 x 10<sup>-9</sup> | 0, 9 x 10<sup>-1</sup> |
| rs7920721 | 10 | 11720308 | ECDH3 | A/G | 0.389 | 1.08 (1.05-1.11) | 1.9 x 10<sup>-7</sup> | 1.07 (1.02-1.12) | 3.2 x 10<sup>-3</sup> | 1.08 (1.05-1.11) | 2.3 x 10<sup>-9</sup> | 0, 8 x 10<sup>-1</sup> |
| Previous genome-wide significant loci not reaching significance |  |  |  |  |  |  |  |  |  |  |  |  |
| rs190982 | 5 | 88223420 | MEF2C | A/G | 0.390 | 0.95 (0.92-0.97) | 2.8 x 10<sup>-4</sup> | 0.93 (0.89-0.98) | 2.7 x 10<sup>-3</sup> | 0.94 (0.92-0.97) | 2.8 x 10<sup>-6</sup> | 0, 6 x 10<sup>-1</sup> |
| rs4723711 | 7 | 37844263 | NME8 | A/T | 0.356 | 0.95 (0.92-0.98) | 2.7 x 10<sup>-4</sup> | 0.91 (0.87-0.95) | 9.5 x 10<sup>-5</sup> | 0.94 (0.91-0.96) | 2.8 x 10<sup>-7</sup> | 0, 5 x 10<sup>-1</sup> |

aVariants showing the best level of association after meta-analysis of stages 1 and 2.
bBuild 37, assembly hg19.
cBased on position of top SNP in reference to the refSeq assembly
dAverage in the discovery sample.
eCalculated with respect to the minor allele.
fCochran's Q test
gPreviously the ZCWPW1 locus.
hPreviously the CELF1 locus.

**Figure 1.**
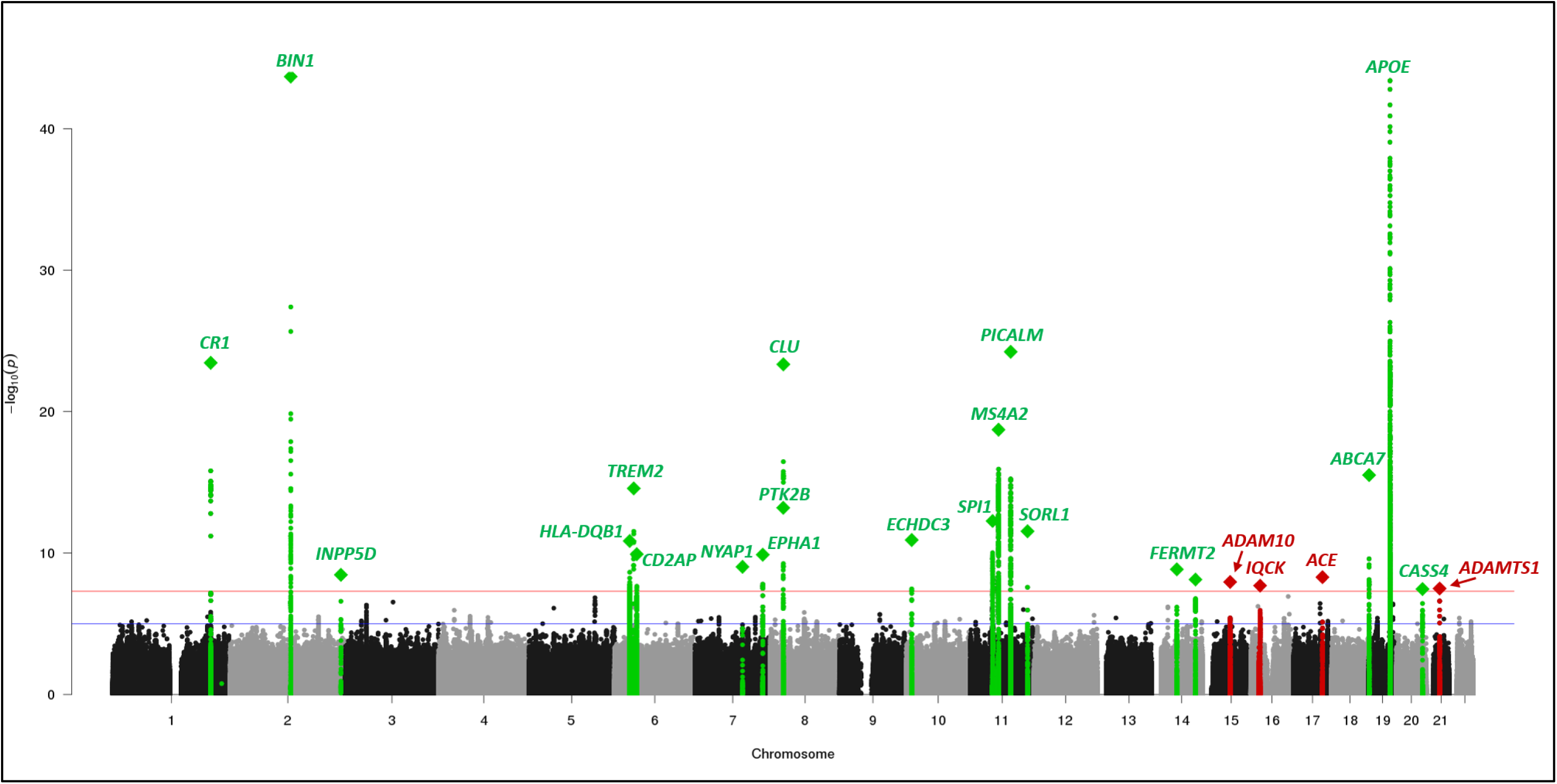
Manhattan plot of meta-analysis of Stage 1, 2 and 3 results for genome-wide association with Alzheimer’s disease. The threshold for genome-wide significance (*P* < 5 × 10^−8^) is indicated by the red line, while the blue line represents the suggestive threshold (*P* < 1 × 10-5). Loci previously identified by the Lambert et al. 2013 IGAP GWAS are shown in green, and newly associated loci are shown in red. Diamonds represent variants with the smallest *P* values for each genome-wide locus.

Stage 3A and meta-analysis of all three stages for these 6 variants (excluding the *TREM2* signal, see **Supplementary Figure 1** for workflow) identified five genome-wide significant sites. In addition to ECDH3, this included four new genome-wide AD risk signals at *IQCK, ADAMTS1, ACE* and *ADAM10* not previously described in other AD GWAS (**Table 2 and Supplementary Figures 3-7**)*. ACE* and *ADAM10* were previously reported as AD candidate genes^14–18^ that were not replicated in some subsequent studies^19–21,17,22^. We also extended the analyses of the two loci (*NME8* and *MEF2C*) in stage 3 that were previously genome-wide significant in our 2013 meta-analysis. These loci were not genome-wide significant in our current study and will deserve further investigations (*NME8*: *P* = 2.8×10^−6^; *MEF2C*: *P* = 2.8×10^−7^). Of note, GCTA-COJO^23^ conditional analysis of the genome-wide loci indicates that *TREM2* and three other loci (*BIN1*, *ABCA7*, and *PTK2B/CLU*) have multiple independent LOAD association signals (**Supplementary Table 5**), suggesting that the genetic variance associated with some GWAS loci is probably under-estimated.

**Table 2.**
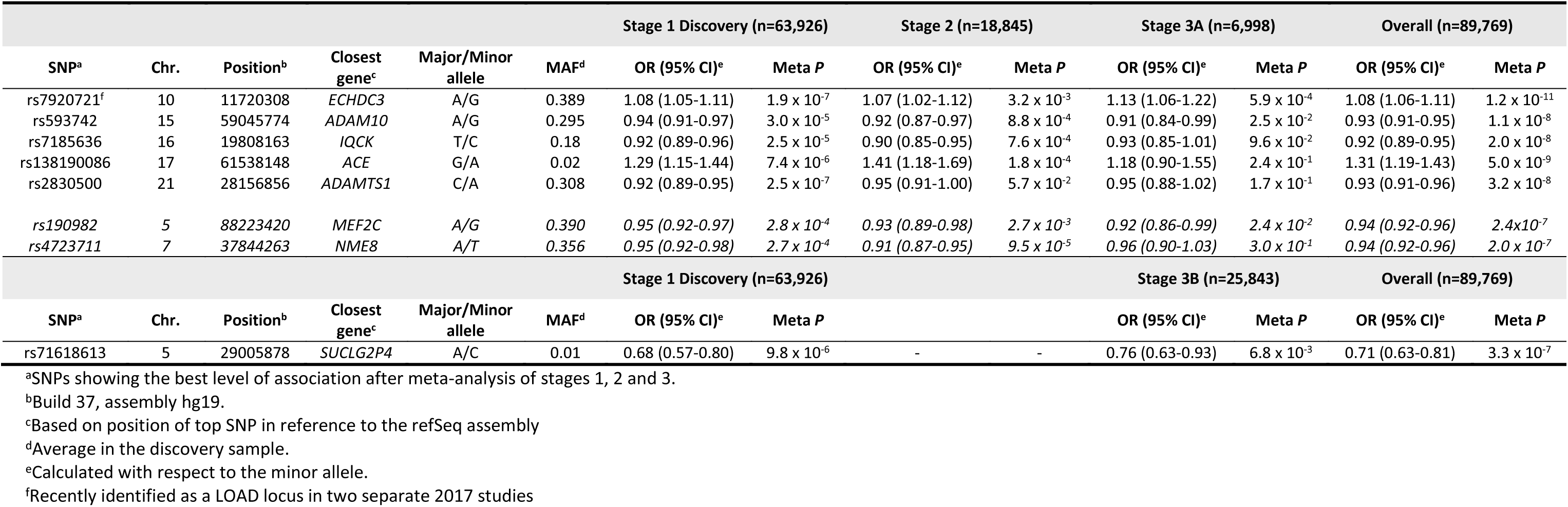
Summary of discovery Stage 1, Stage 2, Stage 3 (A and B), and overall meta-analyses results for potential novel loci reaching P <5.10^−7^.

We also selected 33 SNPs from stage 1 (28 common variants and 5 rare variants in loci not well captured in the I-select chip; see supplementary material and methods section for full selection criteria) for genotyping in stage 3B (including populations of stage 2 and stage 3A). We nominally replicated a rare variant (rs71618613) within an intergenic region near *SUCLG2P4* (MAF = 0.01; *P* = 6.8×10^−3^; combined-*P* = 3.3×10^−7^) and a low-frequency variant in the *TREM2* region (rs114812713, MAF=0.03, P = 1.4×10^−2^; combined-*P* = 4.2×10^−13^) in the gene *OARD1* that may represent an independent signal according to our conditional analysis (**Table 2, Supplementary Figures 8-9, Supplementary Table 5 and 6**).

To evaluate the biological significance of the newly identified signals and those found previously, we pursued four strategies: expression-quantitative trait loci (eQTL) analyses, differential expression in AD versus control brains, gene cluster/pathway analyses, and expression in AD-relevant tissues^24,25^. For the 24 signals reported here, other evidence indicates that *APOE*^26,27^*, ABCA7*^28,29^*, BIN1*^30^*, TREM2*^11,12^*, SORL1*^31,32^*, ADAM10*^33^*, SPI1*^34^, *and CR1*^35^ are the true AD risk gene, though there is a possibility that multiple risk genes exist in these regions^36^. Because many GWAS loci are intergenic, and the closest gene to the sentinel variant may not be the actual risk gene, in these analyses, we considered all genes within ±500kb of the sentinel variant linkage disequilibrium (LD) regions (r^2^ ≥ 0.5) for each locus as a candidate AD gene (**Supplementary Table 7**).

For eQTL analyses, we identified variants in LD with sentinel variants for each locus. For these variants, there were cis-acting eQTLs for 117 genes, with 92 eQTL-controlled genes in AD relevant tissues **(Supplementary Tables 8-11**). For our newly identified loci, the most significant eQTLs for the *ADAM10* signal were for *ADAM10* in blood (*P* = 1.21×10^−13^). For the *IQCK* signal, the top eQTL was for *DEF8* in monocytes (*P* = 5.75×10^−48^). For the *ADAMTS1*, signal, the most significant eQTL was for *ADAMTS1* in blood (*P* = 7.56×10^−7^). No eQTLs were found for the *ACE* locus. These results indicate that *ADAM10*, *ADAMTS1*, and *DEF8* may be the genes responsible for the observed association signal. For previously identified loci, there were eQTLs for *BIN1* in monocytes (*P* = 3.46×10^−67^), *PVRIG* in blood at the *NYAP1* locus (P = 2.02×10^−221^), and *SLC24A4* in monocytes (*P* = 1.27×10^−34^).

To study the differential expression of genes in brains of AD patients versus controls, we used thirteen expression studies^37^. Of 469 protein coding genes within the genome-wide loci, we found 87 upregulated and 55 downregulated genes that were differentially expressed in the same direction in two or more studies. These include four genes at the *ADAM10* locus (*ADAM10* and *SLTM,* each upregulated in two studies; *AQP9*, downregulated in three studies; and *LIPC*, downregulated in two studies), three genes in the *IQCK* locus (*GPRC5B, CCP10*, and *GDE1* upregulated in 13, six and four studies, respectively), six genes in the *ACE* locus (*MAP3K3*, *KCNH6* and *FTSJ3*, upregulated in seven, two and two studies respectively; and *DDX42*, *PSMC5* and *TANC2*, downregulated in seven, five and three studies respectively), and three genes in the *ADAMTS1* locus (*ADAMTS1, CYYR1,* and *ADAMTS5,* upregulated in ten, two and two studies respectively) (**Supplementary Table 12**). For previously described loci, differentially expressed genes included *TFEB near TREM2, MS4A6A* (upregulated in 10 studies) at the chromosome 11 *MS4A* gene cluster, and *FERMT2* (upregulated in 9 studies) on chromosome 14, among others. Brain RNA-seq data reveals many of these differentially expressed candidate genes are expressed in AD-relevant cell types (**Supplementary Table 12**).

We conducted pathway analyses (MAGMA^38^) using five gene set resources. Analysis were conducted separately for common (MAF > 0.01) and rare variants (MAF < 0.01). For common variants, we detected four function clusters including: 1) APP metabolism/Aβ-formation (regulation of beta-amyloid formation: *P* = 4.56×10^−7^ and regulation of amyloid precursor protein catabolic process: *P* = 3.54×10^−6^), 2) tau protein binding (*P* = 3.19×10^−5^), 3) lipid metabolism (four pathways including protein-lipid complex assembly: *P* = 1.45×10^−7^), and 4) immune response (P = 6.32×10^−5^) (**Table 3** and **Supplementary Table 13**). Enrichment of the four pathways remains after removal of genes in the *APOE* region. When *APOE*-region genes and genes in the vicinity of genome-wide significant genes are removed, tau shows moderate association (*P* = 0.027) and lipid metabolism and immune related pathways show strong associations (*P* < 0.001) (**Supplementary Table 14**). Genes driving these enrichments (i.e. having a gene-wide *P* < 0.05) include *SCNA*, a Parkinson’s risk gene that encodes alpha-synuclein, the main component of Lewy bodies, and may play a role in tauopathies^39,40^, for the tau pathway; apolipoprotein genes (*APOM*, *APOA5*) and *ABCA1*, a major regulator of cellular cholesterol, for the lipid metabolism pathways; and 52 immune pathway genes (**Supplementary Table 15**). While no pathways were significantly enriched for rare variants, lipid and Aβ-pathways did have nominal significance in rare-variant-only analyses. Importantly, we also observe a highly significant correlation between common and rare pathway gene results (*P* = 1.32×10^−7^), suggesting that risk AD genes and pathways are enriched for rare variants. In fact, 50 different genes within tau, lipid, immunity and Aβ pathways show nominal association (*P* < 0.05) with LOAD (**Supplementary Table 15**).

**Table 3.** Significant pathways (q-value≤0.05) from MAGMA pathway analysis for common SNV and rare SNV subsets.

| Pathway | N genes in pathway in dataset | Common SNVs *P* * | Common SNVs q-value | Rare SNVs *P* * | Rare SNVs q-value | Pathway description |
| --- | --- | --- | --- | --- | --- | --- |
| GO:65005 | 20 | 1.45E-07* | 9.53E-04 | 6.76E-02 | 8.42E-01 | protein-lipid complex assembly |
| GO:1902003 | 10 | 4.56E-07* | 1.49E-03 | 4.94E-02 | 8.42E-01 | regulation of beta-amyloid formation |
| GO:32994 | 39 | 1.16E-06* | 2.54E-03 | 1.78E-02 | 8.17E-01 | protein-lipid complex |
| GO:1902991 | 12 | 3.54E-06* | 5.80E-03 | 5.66E-02 | 8.42E-01 | regulation of amyloid precursor protein catabolic process |
| GO:43691 | 17 | 5.55E-06* | 6.75E-03 | 3.08E-02 | 8.17E-01 | reverse cholesterol transport |
| GO:71825 | 35 | 6.18E-06* | 6.75E-03 | 1.27E-01 | 8.42E-01 | protein-lipid complex subunit organization |
| GO:34377 | 18 | 1.64E-05* | 1.53E-02 | 1.82E-01 | 8.42E-01 | plasma lipoprotein particle assembly |
| GO:48156 | 10 | 3.19E-05* | 2.61E-02 | 7.77E-01 | 8.54E-01 | tau protein binding |
| GO:2253 | 382 | 6.32E-05* | 4.60E-02 | 2.09E-01 | 8.42E-01 | activation of immune response |

\*Significant after FDR-correction (q-value≤0.05)

To further explore the APP/Aβ-pathway enrichment we analyzed a comprehensive set of 335 APP metabolism genes^41^ curated from the literature. We observed significant enrichment of this gene-set in common variants (*P* = 2.27×10^−4^; *P* = 3.19×10^−4^ excluding *APOE*), with both *ADAM10* and *ACE* nominally significant drivers of this result (**Table 4** and **Supplementary Table 16 and 17**). Several ‘sub-pathways’ were also significantly enriched in the common-variants including ‘clearance and degradation of Aβ’ and ‘aggregation of Aβ’, along with its subcategory ‘microglia’, the latter supporting the recent hypothesis that microglia play a large role in AD^42,43^. Nominal enrichment for risk from rare variants was found for the pathway ‘aggregation of Aβ: chaperone’ and 23 of the 335 genes.

**Table 4.** Top results of pathway analysis of Aβ-beta centered biological network from Campion et al (see Supplementary Table 12 for full results).

| Category | Subcategory | N Genes | Common SNVs P<br>0kb | Common SNVs P 35kb-10kb | Rare SNVs P<br>0kb | Rare SNVs P<br>35kb-10kb |
| --- | --- | --- | --- | --- | --- | --- |
| Aβ -centered biological network (all genes) | -- | 331 | 2.27E-04* | 1.54E-04* | 8.26E-01 | 5.19E-01 |
| Clearance and degradation of Aβ | -- | 74 | 2.18E-04* | 3.27E-03 | 3.13E-01 | 5.11E-01 |
| Clearance and degradation of Aβ | Microglia | 47 | 2.24E-04* | 1.83E-02 | 2.49E-01 | 6.87E-01 |
| Aggregation of Aβ | -- | 35 | 7.09E-04* | 9.93E-03 | 9.02E-02 | 1.68E-01 |
| Aggregation of Aβ | Miscellaneous | 21 | 1.08E-03* | 3.38E-02 | 9.53E-02 | 1.90E-01 |
| APP processing and trafficking | Clathrin/caveolin-dependent endocytosis | 10 | 1.19E-03 | 1.15E-02 | 3.64E-01 | 1.84E-01 |
| Mediator of Aβ toxicity | -- | 51 | 3.82E-02 | 4.69E-02 | 5.89E-01 | 5.70E-01 |
| Mediator of Aβ toxicity | Calcium homeostasis | 6 | 6.90E-02 | 1.21E-01 | 3.96E-01 | 2.54E-01 |
| Mediator of Aβ toxicity | Miscellaneous | 3 | 7.61E-02 | 2.35E-02 | 9.79E-01 | 7.61E-01 |
| Clearance and degradation of Aβ | Enzymatic degradation of Aβ | 15 | 7.77E-02 | 2.63E-02 | 6.10E-01 | 2.95E-01 |
| Mediator of Aβ toxicity | Tau toxicity | 20 | 9.03E-02 | 3.48E-01 | 7.17E-01 | 6.85E-01 |
| Aggregation of Aβ | Chaperone | 9 | 1.52E-01 | 3.09E-01 | 1.98E-01 | 1.13E-02 |

\*Significant after Bonferroni correction for 33 pathway sets tested

To identify candidate genes for our novel loci, we combined results from eQTL, differential expression, AD-relevant tissue expression, and gene function/pathway analyses (**Table 5**). For our *ADAM10* signal, of the 17 genes within this locus, only *ADAM10* meets all our prioritization criteria. In addition, *ADAM10*, the most important α-secretase in the brain, is a component of the non-amyloidogenic pathway of APP metabolism^44^, and sheds *TREM2*^45^, an innate immunity receptor expressed selectively in microglia. Over-expression of *ADAM10* in mouse models can halt Aβ production and subsequent aggregation^46^. Also two rare *ADAM10* mutations segregating with disease in LOAD families increased Aβ plaque load in “Alzheimer-like” mice, with diminished α-secretase activity from the mutations likely the causal mechanism^15,33^. For the *IQCK* signal three of the 12 genes at the locus are potential candidate genes: *IQCK*, *DEF8*, and *GPRC5B.* The latter is a regulator of neurogenesis^47,48^ and inflammatory signalling in obesity^49^. Of the 23 genes in the *ACE* locus, two meet three of the four prioritization criteria, *PSMC5*, a major regulator of major histocompatibility complex^50,51^, and *CD79B,* a B lymphocyte antigen receptor sub-unit. Candidate gene studies previously associate *ACE* variants with AD risk^16,52,18^, including a strong association in the Wadi Ara, an Israeli Arab community with high risk of AD^17^. However, these studies yielded inconsistent results^19^, and our work is the first to report a clear genome-wide association in NHW at this locus. While our analyses did not prioritize *ACE,* it should not be rejected as a candidate gene, as its expression in AD brain tissue is associated with Aβ load and AD severity^53^. Furthermore, CSF levels of the angiotensin-converting enzyme (ACE) are associated with Aβ levels^54^ and LOAD risk^55^, and studies show ACE can inhibit Aβ toxicity and aggregation^56^. Finally, angiotensin II, a product of ACE function mediates a number of neuropathological processes in AD^57^ and is now a target for intervention in phase II clinical trials of AD^58^. Another novel genome-wide locus reported here *ADAMTS1,* is within 665 kb of *APP* on chromosome 21. Of four genes at this locus (*ADAMTS1*, *ADAMTS5*, *CYYR1*, *CYYR1-AS1*), our analyses nominates *ADAMTS1,* as the likely risk gene, though we cannot rule out that this signal is a regulatory element for *APP*. *ADAMTS1* is elevated in Down Syndrome with neurodegeneration and AD^59^ and is a potential neuroprotective gene^60,61,62^, or a neuroinflammatory gene important to microglial response^63^.

**Table 5.**
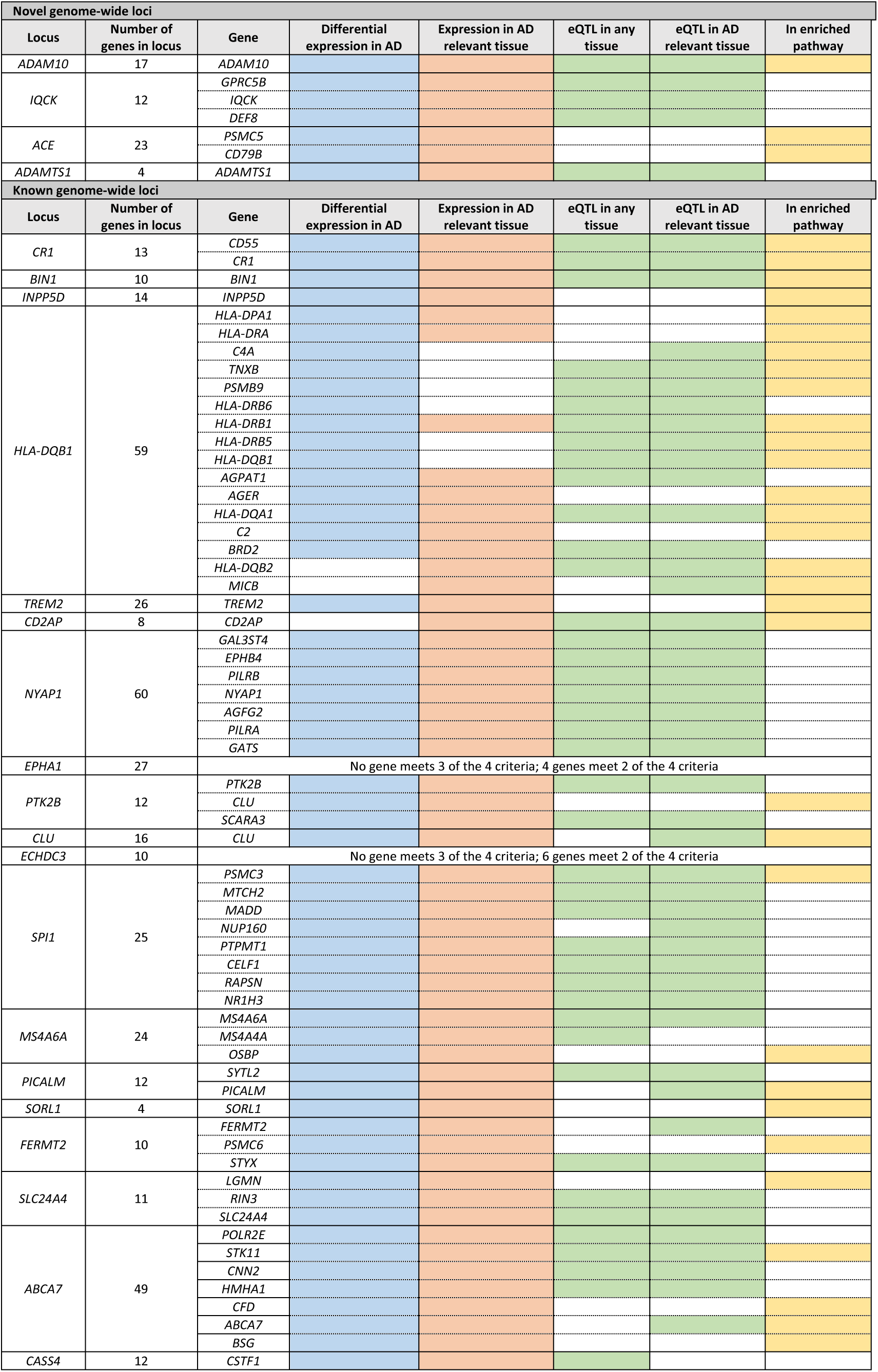
Top prioritized genes in significant loci based on biological evidence. Genes meeting at least 3 of 4 criteria in each locus are listed. The criteria include: 1) differential expression in at least one Alzheimer disease (AD) study, 2) expression in a tissue relevant to AD (astrocytes, neurons, microglia/macrophages, oligodendrocytes), 3) having an eQTL effect on the gene in any tissue, or having an eQTL on the gene in AD relevant tissue, and 4) being involved in a biological pathway enriched in AD (from the current study). Novel genome-wide loci from the current study are listed first, followed by known genome-wide loci.

For previously reported loci, named for the closest gene, applying the same approach for prioritization highlights several genes as described in **Table 5**. It is also interesting to keep in mind that systematic biological screening have also highlighted some of these genes as involved in the APP metabolism (*FERMT2*) or Tau toxicity (*BIN1*, *CD2AP*, *FERMT2*, *CASS4*, *EPHA1*, *PTK2B*)^64–66^. Pathway, tissue and disease traits enrichment analysis supports the utility of our prioritization method, as the 68 prioritized genes are: 1) enriched in substantially more AD relevant pathways and processes, 2) enriched in candidate AD cells such as monocytes (adjusted-*P* = 1.75×10^−6^) and macrophages (adjusted-*P* = 6.46×10^−3^), and 3) increased in associations of dementia-related traits (**Supplementary Table 18 and 19**).

Our work identifies four new genome-wide associations for LOAD and shows that GWAS data combined with high-quality imputation panels can reveal rare disease risk variants (i.e. *TREM2*). The enrichment of rare-variants in pathways associated with AD indicates that additional rare-variants remain to be identified, and larger samples and better imputation panels will facilitate identifying these rare variants. While these rare-variants may not contribute substantially to the predictive value of genetic findings, it will add to the understanding of disease mechanisms and potential drug targets. Discovery of the risk genes at genome-wide loci remains challenging, but we demonstrate that converging evidence from existing and new analyses can prioritize risk genes. We also show that APP metabolism is not only associated with early-onset but also late-onset AD, suggesting that therapies developed by studying early-onset families could also be applicable to the more common late-onset form of the disease. Finally, our analysis showing tau is involved in late-onset AD supports recent evidence that tau may play an early pathological role in AD^67–69^, and confirms that therapies targeting tangle formation/degradation could potentially affect late-onset AD.

## Acknowledgements

**ADGC.** The National Institutes of Health, National Institute on Aging (NIH-NIA) supported this work through the following grants: ADGC, U01 AG032984, RC2 AG036528; Samples from the National Cell Repository for Alzheimer’s Disease (NCRAD), which receives government support under a cooperative agreement grant (U24 AG21886) awarded by the National Institute on Aging (NIA), were used in this study. We thank contributors who collected samples used in this study, as well as patients and their families, whose help and participation made this work possible; Data for this study were prepared, archived, and distributed by the National Institute on Aging Alzheimer’s Disease Data Storage Site (NIAGADS) at the University of Pennsylvania (U24-AG041689-01); NACC, U01 AG016976; NIA LOAD (Columbia University), U24 AG026395, U24 AG026390, R01AG041797; Banner Sun Health Research Institute P30 AG019610; Boston University, P30 AG013846, U01 AG10483, R01 CA129769, R01 MH080295, R01 AG017173, R01 AG025259, R01 AG048927, R01AG33193, R01 AG009029; Columbia University, P50 AG008702, R37 AG015473, R01 AG037212, R01 AG028786; Duke University, P30 AG028377, AG05128; Emory University, AG025688; Group Health Research Institute, UO1 AG006781, UO1 HG004610, UO1 HG006375, U01 HG008657; Indiana University, P30 AG10133, R01 AG009956, RC2 AG036650; Johns Hopkins University, P50 AG005146, R01 AG020688; Massachusetts General Hospital, P50 AG005134; Mayo Clinic, P50 AG016574, R01 AG032990, KL2 RR024151; Mount Sinai School of Medicine, P50 AG005138, P01 AG002219; New York University, P30 AG08051, UL1 RR029893, 5R01AG012101, 5R01AG022374, 5R01AG013616, 1RC2AG036502, 1R01AG035137; North Carolina A&T University, P20 MD000546, R01 AG28786-01A1; Northwestern University, P30 AG013854; Oregon Health & Science University, P30 AG008017, R01 AG026916; Rush University, P30 AG010161, R01 AG019085, R01 AG15819, R01 AG17917, R01 AG030146, R01 AG01101, RC2 AG036650, R01 AG22018; TGen, R01 NS059873; University of Alabama at Birmingham, P50 AG016582; University of Arizona, R01 AG031581; University of California, Davis, P30 AG010129; University of California, Irvine, P50 AG016573; University of California, Los Angeles, P50 AG016570; University of California, San Diego, P50 AG005131; University of California, San Francisco, P50 AG023501, P01 AG019724; University of Kentucky, P30 AG028383, AG05144; University of Michigan, P50 AG008671; University of Pennsylvania, P30 AG010124; University of Pittsburgh, P50 AG005133, AG030653, AG041718, AG07562, AG02365; University of Southern California, P50 AG005142; University of Texas Southwestern, P30 AG012300; University of Miami, R01 AG027944, AG010491, AG027944, AG021547, AG019757; University of Washington, P50 AG005136, R01 AG042437; University of Wisconsin, P50 AG033514; Vanderbilt University, R01 AG019085; and Washington University, P50 AG005681, P01 AG03991, P01 AG026276. The Kathleen Price Bryan Brain Bank at Duke University Medical Center is funded by NINDS grant # NS39764, NIMH MH60451 and by Glaxo Smith Kline. Support was also from the Alzheimer’s Association (LAF, IIRG-08-89720; MP-V, IIRG-05-14147), the US Department of Veterans Affairs Administration, Office of Research and Development, Biomedical Laboratory Research Program, and BrightFocus Foundation (MP-V, A2111048). P.S.G.-H. is supported by Wellcome Trust, Howard Hughes Medical Institute, and the Canadian Institute of Health Research. Genotyping of the TGEN2 cohort was supported by Kronos Science. The TGen series was also funded by NIA grant AG041232 to AJM and MJH, The Banner Alzheimer’s Foundation, The Johnnie B. Byrd Sr. Alzheimer’s Institute, the Medical Research Council, and the state of Arizona and also includes samples from the following sites: Newcastle Brain Tissue Resource (funding via the Medical Research Council, local NHS trusts and Newcastle University), MRC London Brain Bank for Neurodegenerative Diseases (funding via the Medical Research Council),South West Dementia Brain Bank (funding via numerous sources including the Higher Education Funding Council for England (HEFCE), Alzheimer’s Research Trust (ART), BRACE as well as North Bristol NHS Trust Research and Innovation Department and DeNDRoN), The Netherlands Brain Bank (funding via numerous sources including Stichting MS Research, Brain Net Europe, Hersenstichting Nederland Breinbrekend Werk, International Parkinson Fonds, Internationale Stiching Alzheimer Onderzoek), Institut de Neuropatologia, Servei Anatomia Patologica, Universitat de Barcelona. ADNI data collection and sharing was funded by the National Institutes of Health Grant U01 AG024904 and Department of Defense award number W81XWH-12-2-0012. ADNI is funded by the National Institute on Aging, the National Institute of Biomedical Imaging and Bioengineering, and through generous contributions from the following: AbbVie, Alzheimer’s Association; Alzheimer’s Drug Discovery Foundation; Araclon Biotech; BioClinica, Inc.; Biogen; Bristol-Myers Squibb Company; CereSpir, Inc.; Eisai Inc.; Elan Pharmaceuticals, Inc.; Eli Lilly and Company; EuroImmun; F. Hoffmann-La Roche Ltd and its affiliated company Genentech, Inc.; Fujirebio; GE Healthcare; IXICO Ltd.; Janssen Alzheimer Immunotherapy Research & Development, LLC.; Johnson & Johnson Pharmaceutical Research & Development LLC.; Lumosity; Lundbeck; Merck & Co., Inc.; Meso Scale Diagnostics, LLC.; NeuroRx Research; Neurotrack Technologies; Novartis Pharmaceuticals Corporation; Pfizer Inc.; Piramal Imaging; Servier; Takeda Pharmaceutical Company; and Transition Therapeutics. The Canadian Institutes of Health Research is providing funds to support ADNI clinical sites in Canada. Private sector contributions are facilitated by the Foundation for the National Institutes of Health (www.fnih.org). The grantee organization is the Northern California Institute for Research and Education, and the study is coordinated by the Alzheimer’s Disease Cooperative Study at the University of California, San Diego. ADNI data are disseminated by the Laboratory for Neuro Imaging at the University of Southern California. We thank Drs. D. Stephen Snyder and Marilyn Miller from NIA who are *ex-officio* ADGC members.

**EADI.** This work has been developed and supported by the LABEX (laboratory of excellence program investment for the future) DISTALZ grant (Development of Innovative Strategies for a Transdisciplinary approach to ALZheimer’s disease) including funding from MEL (Metropole européenne de Lille), ERDF (European Regional Development Fund) and Conseil Régional Nord Pas de Calais. This work was supported by INSERM, the National Foundation for Alzheimer’s disease and related disorders, the Institut Pasteur de Lille and the Centre National de Génotypage, the JPND PERADES and the FP7 AgedBrainSysBio. The Three-City Study was performed as part of collaboration between the Institut National de la Santé et de la Recherche Médicale (Inserm), the Victor Segalen Bordeaux II University and Sanofi-Synthélabo. The Fondation pour la Recherche Médicale funded the preparation and initiation of the study. The 3C Study was also funded by the Caisse Nationale Maladie des Travailleurs Salariés, Direction Générale de la Santé, MGEN, Institut de la Longévité, Agence Française de Sécurité Sanitaire des Produits de Santé, the Aquitaine and Bourgogne Regional Councils, Agence Nationale de la Recherche, ANR supported the COGINUT and COVADIS projects. Fondation de France and the joint French Ministry of Research/INSERM “Cohortes et collections de données biologiques” programme. Lille Génopôle received an unconditional grant from Eisai. The Three-city biological bank was developed and maintained by the laboratory for genomic analysis LAG-BRC - Institut Pasteur de Lille.

*Belgium samples*: Research at the Antwerp site is funded in part by the Interuniversity Attraction Poles program of the Belgian Science Policy Office, the Foundation for Alzheimer Research (SAO-FRA), a Methusalem Excellence Grant of the Flemish Government, the Research Foundation Flanders (FWO), the Special Research Fund of the University of Antwerp, Belgium. KB is a postdoctoral fellow of the FWO. The Antwerp site authors thank the personnel of the VIB Genetic Service Facility, the Biobank of the Institute Born-Bunge and the Departments of Neurology and Memory Clinics at the Hospital Network Antwerp and the University Hospitals Leuven.

*Finish sample collection*: Financial support for this project was provided by Academy of Finland (grant number 307866), Sigrid Jusélius Foundation and the Strategic Neuroscience Funding of the University of Eastern Finland

*Swedish sample collection*: Financially supported in part by the Swedish Brain Power network, the Marianne and Marcus Wallenberg Foundation, the Swedish Research Council (521-2010-3134, 2015-02926), the King Gustaf V and Queen Victoria’s Foundation of Freemasons, the Regional Agreement on Medical Training and Clinical Research (ALF) between Stockholm County Council and the Karolinska Institutet, the Swedish Brain Foundation and the Swedish Alzheimer Foundation”.

**CHARGE.** Infrastructure for the CHARGE Consortium is supported in part by National Heart, Lung, and Blood Institute grant HL105756 (Psaty) and RC2HL102419 (Boerwinkle) and the neurology working group by grants from the National Institute on Aging, R01 AG033193, U01 AG049505 and U01AG52409.

*Rotterdam (RS)*. The Rotterdam Study is funded by Erasmus Medical Center and Erasmus University, Rotterdam, the Netherlands Organization for Health Research and Development (ZonMw), the Research Institute for Diseases in the Elderly (RIDE), the Ministry of Education, Culture and Science, the Ministry for Health, Welfare and Sports, the European Commission (DG XII), and the municipality of Rotterdam. The authors are grateful to the study participants, the staff from the Rotterdam Study, and the participating general practitioners and pharmacists. Generation and management of GWAS genotype data for the Rotterdam Study (RS-I, RS-II, RS-III) was executed by the Human Genotyping Facility of the Genetic Laboratory of the Department of Internal Medicine, Erasmus MC, Rotterdam, the Netherlands. The GWAS data sets are supported by the Netherlands Organization of Scientific Research NWO Investments (175.010.2005.011, 911-03-012), the Genetic Laboratory of the Department of Internal Medicine, Erasmus MC, the Research Institute for Diseases in the Elderly (014-93-015; RIDE2), and the Netherlands Genomics Initiative (NGI)/Netherlands Organization for Scientific Research (NWO) Netherlands Consortium for Healthy Aging (NCHA), project 050-060-810. This study makes use of an extended data set of RS-II and RS-III samples based on Illumina Omni 2.5 and 5.0 GWAS genotype data. This data set was funded by the Genetic Laboratory of the Department of Internal Medicine, the Department of Forensic Molecular Biology, and the Department of Dermatology, Erasmus MC, Rotterdam, the Netherlands. We thank M. Jhamai, M. Verkerk, L. Herrera, M. Peters, and C. Medina-Gomez for their help in creating the GWAS database. The work for this manuscript was further supported by ADAPTED: Alzheimer’s Disease Apolipoprotein Pathology for Treatment Elucidation and Development (115975); the CoSTREAM project (http://www.costream.eu/); and funding from the European Union’s Horizon 2020 research and innovation programme under grant agreement 667375. The ERF study as a part of EUROSPAN (European Special Populations Research Network) was supported by European Commission FP6 STRP grant number 018947 (LSHG-CT-2006-01947) and also received funding from the European Community’s Seventh Framework Programme (FP7/2007-2013)/grant agreement HEALTH-F4-2007-201413 by the European Commission under the programme "Quality of Life and Management of the Living Resources" of 5th Framework Programme (no. QLG2-CT-2002-01254). High-throughput analysis of the ERF data was supported by a joint grant from the Netherlands Organization for Scientific Research and the Russian Foundation for Basic Research (NWO-RFBR 047.017.043).

*AGES*. The AGES study has been funded by NIA contracts N01-AG-12100 and HHSN271201200022C with contributions from NEI, NIDCD, and NHLBI, the NIA Intramural Research Program, Hjartavernd (the Icelandic Heart Association), and the Althingi (the Icelandic Parliament).

*Cardiovascular Health Study (CHS)*. This research was supported by contracts HHSN268201200036C, HHSN268200800007C, N01HC55222, N01HC85079, N01HC85080, N01HC85081, N01HC85082, N01HC85083, and N01HC85086 and grant U01HL080295 *and* U01HL130114 from the National Heart, Lung, and Blood Institute (NHLBI), with additional contribution from the National Institute of Neurological Disorders and Stroke (NINDS). Additional support was provided by R01AG033193, R01AG023629, R01AG15928, and R01AG20098 and by U01AG049505 from the National Institute on Aging (NIA). The provision of genotyping data was supported in part by the National Center for Advancing Translational Sciences, CTSI grant UL1TR000124, and National Institute of Diabetes and Digestive and Kidney Disease Diabetes Research Center (DRC) grant DK063491 to the Southern California Diabetes Endocrinology Research Center. A full list of CHS principal investigators and institutions can be found at https://chs-nhlbi.org/. The content is solely the responsibility of the authors and does not necessarily represent the official views of the US National Institutes of Health.

*Framingham Heart Study*. This work was supported by the National Heart, Lung, and Blood Institute’s Framingham Heart Study (contracts N01-HC-25195 and HHSN268201500001I). This study was also supported by grants from the National Institute on Aging: R01AG033193, U01AG049505, U01AG52409, R01AG054076 (S. Seshadri). S. Seshadri and A.L.D. were also supported by additional grants from the National Institute on Aging (R01AG049607, R01AG033040) and the National Institute of Neurological Disorders and Stroke (R01-NS017950, NS100605). The content is solely the responsibility of the authors and does not necessarily represent the official views of the US National Institutes of Health.

*Fundació ACE*. We would like to thank patients and controls who participated in this project. We also want to thank the private sponsors supporting the basic and clinical projects of our institution (Grifols SA, Piramal AG, Laboratorios Echevarne, Araclon Biotech S.A. and Fundació ACE). We are indebted to Trinitat Port-Carbó legacy and her family for their support of Fundació ACE research programs. Fundació ACE collaborates with the Centro de Investigación Biomédica en Red sobre Enfermedades Neurodegenerativas (CIBERNED, Spain) and is one of the participating centers of the Dementia Genetics Spanish Consortium (DEGESCO). A.R. and M.B. are receiving support from the European Union/EFPIA Innovative Medicines Initiative Joint Undertaking ADAPTED and MOPEAD projects (Grants No. 115975 and 115985 respectively). M.B. and A.R. are also supported by national grants PI13/02434, PI16/01861 and PI17/01474. Acción Estratégica en Salud integrated in the Spanish National R + D + I Plan and financed by ISCIII (Instituto de Salud Carlos III)-Subdirección General de Evaluación and the Fondo Europeo de Desarrollo Regional (FEDER- “Una manera de Hacer Europa”). Genome Resesarch @ Fundació ACE project (GR@ACE) is supported by Fundación bancaria “La Caixa”, ISCIII and Grifols SA.

**GERAD/PERADES.** We thank all individuals who participated in this study. Cardiff University was supported by the Wellcome Trust, Alzheimer’s Society (AS; grant RF014/164), the Medical Research Council (MRC; grants G0801418/1, MR/K013041/1, MR/L023784/1), the European Joint Programme for Neurodegenerative Disease (JPND, grant MR/L501517/1), Alzheimer’s Research UK (ARUK, grant ARUK-PG2014-1), Welsh Assembly Government (grant SGR544:CADR), a donation from the Moondance Charitable Foundation, and the UK Dementia Research Institute at Cardiff. Cambridge University acknowledges support from the MRC. ARUK supported sample collections at the Kings College London, the South West Dementia Bank, Universities of Cambridge, Nottingham, Manchester and Belfast. King’s College London was supported by the NIHR Biomedical Research Centre for Mental Health and Biomedical Research Unit for Dementia at the South London and Maudsley NHS Foundation Trust and Kings College London and the MRC. Alzheimer’s Research UK (ARUK) and the Big Lottery Fund provided support to Nottingham University. Ulster Garden Villages, AS, ARUK, American Federation for Aging Research, NI R&D Office and the Royal College of Physicians/Dunhill Medical Trust provided support for Queen’s University, Belfast. The University of Southampton acknowledges support from the AS. The MRC and Mercer’s Institute for Research on Ageing supported the Trinity College group. DCR is a Wellcome Trust Principal Research fellow. The South West Dementia Brain Bank acknowledges support from Bristol Research into Alzheimer’s and Care of the Elderly. The Charles Wolfson Charitable Trust supported the OPTIMA group. Washington University was funded by NIH grants, Barnes Jewish Foundation and the Charles and Joanne Knight Alzheimer’s Research Initiative. Patient recruitment for the MRC Prion Unit/UCL Department of Neurodegenerative Disease collection was supported by the UCLH/UCL Biomedical Centre and their work was supported by the NIHR Queen Square Dementia BRU. LASER-AD was funded by Lundbeck SA. The Bonn group would like to thank Dr. Heike Koelsch for her scientific support. The Bonn group was funded by the German Federal Ministry of Education and Research (BMBF): Competence Network Dementia (CND) grant number 01GI0102, 01GI0711, 01GI0420. The AgeCoDe study group was supported by the German Federal Ministry for Education and Research grants 01 GI 0710, 01 GI 0712, 01 GI 0713, 01 GI 0714, 01 GI 0715, 01 GI 0716, 01 GI 0717. Genotyping of the Bonn case-control sample was funded by the German centre for Neurodegenerative Diseases (DZNE), Germany. The GERAD Consortium also used samples ascertained by the NIMH AD Genetics Initiative. HH was supported by a grant of the Katharina-Hardt-Foundation, Bad Homburg vor der Höhe, Germany. The KORA F4 studies were financed by Helmholtz Zentrum München; German Research Center for Environmental Health; BMBF; German National Genome Research Network and the Munich Center of Health Sciences. The Heinz Nixdorf Recall cohort was funded by the Heinz Nixdorf Foundation (Dr. Jur. G.Schmidt, Chairman) and BMBF. Coriell Cell Repositories is supported by NINDS and the Intramural Research Program of the National Institute on Aging. We acknowledge use of genotype data from the 1958 Birth Cohort collection, funded by the MRC and the Wellcome Trust which was genotyped by the Wellcome Trust Case Control Consortium and the Type-1 Diabetes Genetics Consortium, sponsored by the National Institute of Diabetes and Digestive and Kidney Diseases, National Institute of Allergy and Infectious Diseases, National Human Genome Research Institute, National Institute of Child Health and Human Development and Juvenile Diabetes Research Foundation International. The Bonn samples are part of the German Dementia Competance Network (DCN) and the German Research Network on Degenerative Dementia (KNDD), which are funded by the German Federal Ministry of Education and Research (grants KND: 01G10102, 01GI0420, 01GI0422, 01GI0423, 01GI0429, 01GI0431, 01GI0433, 04GI0434; grants KNDD: 01GI1007A, 01GI0710, 01GI0711, 01GI0712, 01GI0713, 01GI0714, 01GI0715, 01GI0716, 01ET1006B). Markus M Nothen is a member of the German Research Foundation (DFG) cluster of excellence ImmunoSensation. Funding for Saarland University was provided by the German Federal Ministry of Education and Research (BMBF), grant number 01GS08125 to Matthias Riemenschneider. The University of Washington was supported by grants from the National Institutes of Health (R01-NS085419 and R01-AG044546), the Alzheimer’s Association (NIRG-11-200110) and the American Federation for Aging Research (Carlos Cruchaga was recipient of a New Investigator Award in Alzheimer’s disease). Brigham Young University was supported by the Alzheimer’s Association (MNIRG-11-205368), the BYU Gerontology Program and the National Institutes of Health (R01-AG11380, R01-AG021136, P30-S069329-01, R01-AG042611). We also acknowledge funding from the Institute of Neurology, UCL, London who were supported in part by the ARUK via an anonymous donor, and by a fellowship to Dr Guerreiro. Seripa, Urbano and Masullo’s participation in the study was completely supported by Ministerodella Salute”, I.R.C.C.S. Research Program, Ricerca Corrente 2015-2017, Linea n. 2 “Malattiecomplesse e terapie innovative” and by the “5 × 1000” voluntary contribution. AddNeuromed is supported by InnoMed, an Integrated Project funded by the European Union Sixth Framework programme priority FP6-2004-LIFESCIHEALTH-5, Life Sciences, Genomics and Biotechnology for Health. We are grateful to the Wellcome Trust for awarding a Principal Research Fellowship to Rubensztein (095317/Z/11/Z). Matthias Riemenschneider was funded by the BMBF NGFN Grant 01GS08125. BN supported by FondazioneCassa di Risparmio di Pistoia e Pescia (grants 2014.0365, 2011.0264 and 2013.0347). Harald Hampel is supported by the AXA Research Fund, the Fondation Universite Pierre et Marie Curie and the “Fondation pour la Recherchesur Alzheimer”, Paris, France. The research leading to these results has received funding from the program “Investissementsd’ avenir” ANR-10-IAIHU-06 (Agence Nationale de la Recherche-10-IA Agence Institut Hospitalo-Universitaire-6. The Santa Lucia Foundation and the Fondazione Ca’ Granda IRCCS Ospedale Policlinico, Italy, acknowledge the Italian Ministry of Health (grant RC 10.11.12.13/A). We acknowledge Maria A Pastor (Department of Neurology, University of Navarra Medical School and Neuroimaging Laboratory, Center for Applied Medical Research, Pamplona, Spain), for providing DNA samples.

## Author Contributions

### ADGC

*Study design or conception:* A.C.N., A.A.-W., E.R.M., K.H.-N., A.B.K., B.N.V., G.W.B., O.V., M.Butkiewics, W.B., Y.Song, G.D.S., M.A.P.-V. *Sample contribution:* S.S.M., P.K.C., R.B., P.M.A., M.S.A., D. Beekly, D. Blacker, R.S. Doody, T.J.F., M.P.F., B.Ghetti, R.M.H., M.I.K., M.J.K., C.K., W.K., E.B.L., R.B.L., T.J.M., R.C.P., E.M.R., J.S.R., D.R.R., M. Sano, P.S.G.-H., D.W.T., C.K.W., R.L.A., L.G.A., S.E.A., S.A., C.S.A., C.T.B., L.L.B., S. Barral, T.G.B., J.T.B., E.B., T.D.B., B.F.B., J.D.B., A.Boxer, J.R.B., J.M.B., J.D.Buxbaum, N.J.C., C. Cao, C.S.C., C.M.C., R.M.C., M.M.C., H.C.C., D.H.C., E.A.C., C.DeCarli, M.Dick, R.D., N.R.G.-R., D.A.E., K.M.F., K.B.F., D.W.F., M.R.F., S.F., T.M.F., D.R.G., M.Gearing, D.H.G., J.R.G., R.C.G., J.H.G., R.H., L.E.H., L.S.H., M.J.H., C.M.H., B.T.H., G.P.J., E.A., L.W.J., G.J., A. Karydas, J.A.K., R.K., N.W.K., J.H.K., F.M.L., J.J.L., J.B.L., A.I.L., A.P.L., K.L.L., C.G.L., D.C.M., F.M., D.C.Mash, E.M., W.C.M., S.M.M., A.N.M., A.C.M., M.M., B.L.M., C.A.M., J.W.M., J.C.M., A.J.M., S.O., J.M.O., J.E.P., H.L.P., W.P., E.P., A.P., W.W.P., H.P., J.F.Q., A.Raj, M.R., B.R., C.R., J.M.R., E.D.R., E.R., H.J.R., R.N.R., M.A.S., A.J.S., M.L.C., J. Vance, J.A.S., L.S.S., S.Slifer, W.W.S., A.G.S., J.A.Sonnen, S. Spina, R.A.S., R.H.S., R.E.T., J.Q.T., J.C.T., V.M.V.D., L.J.V.E., H.V.V., J.P.V., S.W., K.A.W.-B., K.C.W., J.Williamson, T.S.W., R.L.W., C.B.W., C.-E.Y., L.Y., D.B., P.L.D.J., S.McDonough, C.Cruchaga, A.M.G., N.E.-T., S.G.Y., D.W.D., H.H., L.A.F., J.Haines, R.Mayeux, L.-S.W., G.D.S., M.A.P.-V. *Data generation:* B.W.K., K.H.-N., A.B.K., O.V., L.Q., Y.Z., J. Malamon, B.Dombroski, P.W., L.B.C., M.A., J.R.G., L.-S.W. *Analysis:* B.W.K., A.C.N., A.A.-W., E.R.M., K.H.-N., A.B.K., B.N.V., G.W.B., O.V., M.Butkiewics, W.B., Y.Song, G.D.S., M.A.P.-V. *Manuscript preparation:* B.W.K., G.D.S., M.A.P.-V. *Study supervision/management:* B.W.K., L.A.F., J.Haines, R.Mayeux, L.-S.W., G.D.S., M.A.P.-V.

### EADI

*Study design or conception:* P.A., J.-C.L. *Sample contribution:* K.S., M.Hiltunen, J.E., M.D.Z., I.M., F.S.G., M.C.D.N., D.Wallon, S.E., R.V., P.D.D., A.Squassina, E.R.-R., C.M.F., Y.A.B., H.T., V.Giedraitis, L.Kilander, R.Brundin, L.C., S.Helisalmi, A.M.K., A.Haapasalo, V.S., V.Frisardi, V.Deramecourt, N.F., O.H., C.Dufouil, A.Brice, K.R., B.D., H.Soininen, L.Fratiglioni, L.K., F.Panza, D.H., P.C., L.Lannfelt, F.P., M.Ingelsson, C.G., P.S.-J., C.Berr, J.-F.D., D.C., C.V.B.,J.-F.Deleuze, P.A., J.-C.L. *Data generation:* R.Aloso, J.-G.G., M.-L.M., D.Bacq, F.G., B.F., S.Meslage *Analysis:* B.G.-B., A.Boland, C.Bellenguez *Manuscript preparation:* B.G.-B., P.A., J.-C.L. *Study supervision/management:* P.A., J.-C.L.

### GERAD/PERADES

*Study design or conception:* R.Sims, M.C.O., M.J.O., A.R., P.A.H., J.W. *Sample contribution:* R.Raybould, T.Morgan, P.Hoffman, D.Harold, N.D., N.C.F., J.T.H., Y.P., M.Daniilidou, J.U., D.Galimberti, E.Scarpini, J.Kornhuber, S.P., M.Mayhaus, W.G., A.M.H., S.Lovestone, R.Sussams, C.Holmes, W.M., A.Kawalia, S.Moebus, J.Turton, J.Lord, I.K., A.L., B.L., M.Gill, S.O.-C., A.Ciaramella, C.Caltagirone, C.Cupidi, R.G.M., R.Cecchetti, M.T., D.Craig, D.A., A.G., M.K., O.G., D.Markina, H.Hampel, D.C.R., L.F., B.G., J.J., P.Passmore, J.M.S., J.D.W., M.K.L., P.Proitsi, J.Powell, J.S.K.K., M.Mancuso, U.B., A.Q., G.Livingston, N.J.B., J.Hardy, J.B., R.Guerreiro, E.F., C.Masullo, G.B., L.M., A.H., M.Scherer, M.Reimenschneider, R.Heun, M.Leber, I.H., I.G., M.Hull, J.M., K.Mayo, T.F., D.Drichel, T.D.C., P.Hollingworth, R.Marshall, A.Meggy, G.M., G.L., D.G., G.R., F.J., B.V., E.V., K.-H.J., M.Dichgans, D.Mann, S.P.-B., N.K., H.W., K.M., K.Brown, C.Medway, M.M.N., N.M.H., A.Daniele, A.Bayer, J.G., H.V.D.B., C.Brayne, S.R.-H., A.A.-C., C.E.S., J.Wiltfang, E.C., V.A., A.B.S., J.C., S.M., M.Rossor, N.R., B.N., S.Sorbi, E.S., G.S., P.B., R.C., A.D.S., D.W., G.W., A.C.B., M.G., Y.B.-S., P.M., P.P., V.B., N.W., P.D., R.G., P.G.K., S.L., C.C., J.T., R.Munger, A.R., J.W. *Data generation:* R.Sims, R.Raybould, T.Morgan, P.Hoffman, D.Harold, A.Gerrish, N.D., P.Hollingworth, R.Marshall, A.Meggy, A.R., J.W. *Analysis:* R.Sims, M.V., A.F., N.Badarinarayan, D.Harold, G.M., G.L., D.G., V.E.-P., A.R., J.W. *Manuscript preparation:* R.Sims, T.D.C., P.A.H., J.W. *Study supervision/management:* R.Sims, L.J., V.E.-P., A.R., P.A.H., J.W.

### CHARGE

*Study design or conception:* A.L.D., C.M.V.D., S.S. *Sample contribution:* J.C.B., A.Ruiz, A.L.F., G.E., J.J.H., A.O., M.E.G., H.L., H.Comic, G.Roschupkin, S.Li, I.Hernández, Q.Y., A.S.B., L.T., T.H.M., WT.L., F.R., E.Boerwinkle, J.I.R., A.G.U., S.M.-G., O.L.L., M.B., M.F., N.A., L.J.L., M.A.I., H.S., R.S., V.G., B.M.P. *Data generation:* J.C.B., J.Jakobsdottir, A.Ruiz, A.V.S., X.J., S.-H.C., H.H.A., J.A.B., T.A., E.H., C.Sarnowski, D.V., L.A.C. *Analysis:* J.C.B., S.J.v.d.L., V.C., J.Jakobsdottir, Y.C., S.Ahmad, A.Ruiz, A.V.S., C.C.W., C.M.V.D., S.S. *Manuscript preparation:* S.J.v.d.L., A.Ruiz, B.M.P., C.M.V.D., S.S. *Study supervision/management:* C.M.V.D., S.S.

## Methods

### Samples

All stage I meta-analysis samples are from four Consortia: the Alzheimer’s Disease Genetics Consortium (ADGC), the Cohorts for Heart and Aging Research in Genomic Epidemiology (CHARGE) Consortium, the European Alzheimer’s Disease Initiative (EADI), and the Genetic and Environmental Risk in Alzheimer’s Disease (GERAD) Consortium. Summary demographics of all 37 case-control studies from the four consortia are described in **Table 1** and **Supplementary Tables 1 and 2**. Written informed consent was obtained from study participants or, for those with substantial cognitive impairment, from a caregiver, legal guardian or other proxy. Study protocols for all cohorts were reviewed and approved by the appropriate institutional review boards. Further details of all cohorts can be found in the **Supplementary Note**.

### Pre-imputation genotype chip quality control

Standard quality control (QC) was performed on all datasets individually, including exclusion of individuals with low call rate (<90%), individuals with a high degree of relatedness (pi_hat > 0.98) and variants with low call rate (<95%). Individuals with non-European ancestry according to principal components (PCs) analysis of ancestry informative markers were excluded from the further analysis.

### Imputation and pre-analysis quality control

Following genotype chip QC, each dataset was phased and imputed with data to the 1000 Genomes Project (phase 1 integrated release 3, March 2012)^1^ using SHAPEIT/IMPUTE2^2,3^ or MaCH/Minimac^4,5^ software (**Supplementary Table 3**). All reference population haplotypes were used for the imputation as this method improves accuracy of imputation for low-frequency variants^6^. Common variants (MAF ≥ 0.01%) with an r^2^ < 0.30 from MaCH or an information measure < 0.40 from IMPUTE2 were excluded from further analyses. Rare variants (MAF < 0.01%) with a ‘global’ weighted imputation quality score of < 0.70 were also excluded from analyses. This score was calculated by weighting each variants MACH/IMPUTE2 imputation quality score by study sample size and combining these weighted scores for use as a post-analysis filter. We also required the presence of each variant in 30% of AD cases and 30% of controls across all datasets.

### Association Analysis

The Stage 1 discovery meta-analysis was followed by Stage 2, and Stage 3 (A and B) replication analyses. Stage 2 was data from a custom array with 11,632 assays selected as variants with *P* < 10^−3^ from our 2013 work^7^. Genotypes were determined for 8,362 cases and 10,484 controls (**Supplementary Table 20**). Stage 3A was conducted for variants selected as novel loci from meta-analyses of Stages 1 and 2 with *P* < 5 × 10^−7^ (6 variants) and variants that were previously significant (*P* < 5 × 10^−8^) that were not genome-wide significant after Stages 1 and 2 (2 variants) (3,348 cases and 3,650 controls) (**Supplementary Table 21**). Stage 3B, which combined samples from Stage 2 and 3A, was conducted for variants with MAF < 0.05 and *P* < 1 × 10^−5^ or variants with MAF ≥ 0.05 and *P* < 5 × 10^−6^ from genome regions not covered on the Stage 2 custom array (11,710 cases and 14,133 controls) (**Supplementary Table 6**). For Stages 1, 2, and 3, samples did not overlap.

Stage 1 single variant-based association analysis was conducted on genotype dosages modeling for an additive genotype model and adjusting for age (defined as age-at-onset for cases and age-at-last exam for controls), sex and population substructure using PCs^8^. The score test was implemented on all case-control datasets. This test was shown to be optimal for meta-analysis of rare variants due to its balance between power and control of type 1 error^9^. Family datasets were tested using the R package GWAF^10^, with generalized estimating equations (GEE) implemented for common variants (MAF ≥ 0.01), and a general linear mixed effects model (GLMM) implemented for rare variants (MAF < 0.01), per internal data showing behavior of test statistics for GEE was fine for common variants but inflated for rare variants, while GLMM controlled this rare variant inflation. Variants with regression coefficient |β| > 5 or P value equal to 0 or 1 were excluded from further analysis.

Within-study results for Stage 1 were meta-analyzed in METAL^11^ using an inverse-variance based model with genomic control. The meta-analysis was split into two separate analyses based on the study sample size, with all studies being included in the analysis of common variants (MAF ≥ 0.01), and only studies with a total sample size of 400 or greater being included in the rare variant (MAF < 0.01) analysis. We also conducted a second meta-analysis in METAL using a sample-size weighted meta-analysis model. Results of this model were compared to the inverse-variance weighted meta-analysis, and results that differed by more than 3 logs on both *P*-values were removed from further analysis. Regression coefficients for rare variants can at times be unstable^12^, and this step attempted to control for these problematic variants by using a second method of meta-analysis that may be less sensitive to certain properties of rare variant analysis. In total, 11 variants were removed through this comparison, and most results showed very little difference in *P*-values between the two methods. An additional 106 variants with high heterogeneity between studies (defined as *I*^*2*^ > 75) were removed. Figures for association signals were generated with LocusZoom software^13^. Genome-wide summary statistics are available from The National Institute on Aging Genetics of Alzheimer’s Disease (NIAGADS) website (https://www.niagads.org/). These analyses were conducted by two independent consortia (ADGC and EADI) and then cross-validated. Analyses for Stage 2 and Stage 3 followed these same analysis procedures, except covariate adjustments per cohort, where all analyses were adjusted on sex and age apart from Italian and Swedish cohorts, which were also adjusted for PCs.

GCTA^14^ was used to conduct conditional analysis using 37,635 individuals from the ADGC as a reference panel for calculation of linkage disequilibrium (LD). LDLink^15^ was used to conduct LD, using all 5 CEU populations as the reference for calculations.

### Stage 2 and 3 Genotyping and Quality Control

Datasets for Stage 2 analysis were obtained from previous genotyping from Lambert et al. 2013^7^ of 11,632 single nucleotide variants genotyped using Illumina iSelect technology. Eight variants from Stage 3A were genotyped using Taqman technology. Stage 3B included 23 variants included as part of Sequenom MassArray iPLEX panels and 10 additional variants genotyped using Taqman technology.

Per sample quality checks for genetic sex and relatedness were performed in PLINK. Individuals not matching their reported sex or showing a high degree of relatedness (IBD value of 0.98 or greater) were removed from the analysis. A panel of ancestry-informative markers (AIMs), was used to perform PCA analysis with SMARTPCA from EIGENSOFT 4.2 software^16^, and individuals with non-European ancestry were excluded. Variant quality control was also performed separately in each country including removal of variants missing in more than 10% of individuals, having a Hardy-Weinberg P value in controls lower than 1 × 10^−6^, or a P value for missingness between cases and controls lower than 1 × 10^−6^. Please see Lambert et al. for a more detailed description of the QC procedures followed in Stage 2 analysis. After quality control, 18,845 individuals (8,362 cases and 10,483 controls) were available for the stage 2 analysis. The same quality control measures were applied to data for the Stage 3 variants attained from follow-up genotyping.

### Selection of variants for 3B follow-up genotyping

In order to prioritize variants for genotyping in Stage 3B, we first selected all MAF < 0.05 variants with *P* < 1 × 10^−5^ or MAF ≥ 0.05 variants with *P* < 5 × 10^−6^ in novel loci not covered in the iSelect genotyping from Stage 2 of Lambert et al.^7^ A total of 180 variants were considered for follow up due to meeting the *P*-value criteria and not being in an IGAP 2013 locus. 88 of these variants were in a region covered in the replication genotyping chip from 2013 and thus were removed from further consideration. 33 loci remained after their removal, with 19 loci having only one prioritized variant, which we selected for genotyping. Remaining variants in 14 regions with multiple prioritized variants were then annotated with GWAVA^17^ and CADD^18^ scores (using ANNOVAR^19^), Ensembl Variant Effect Predictor (VEP) Consequences (using Ensembl VEP^20^), GWAS3D^21^, RegulomeDB^22^, and FANTOM5^23^ (using NIAGADS GenomicsDB) in order to rank their functional potential. A CADD score > 10, GWAVA score > 0.5, FATHHM > 0.5, RegulomeDB score < 5 and GWAS3D top p-value score were considered ‘functional’ in the ranking. The top ranked variant for functional potential for each locus with multiple variants was selected for further genotyping and analysis. Removal of 59 variants in regions with multiple variants left 33 total variants for follow-up genotyping.

### Characterization of gene(s) and non-coding features in associated loci

We determined the basepair (bp) boundaries of the search space for potential gene(s) and non-coding features in each of the 24 associated loci (excluding *APOE*) using the ‘proxy search’ mechanism in LDLink^15^. LDLink uses 1000 genomes genotypes to calculate LD for a selected population; in our case all five European population were selected (CEU, TSI, FIN, GBR, and IBS). The boundaries for all variants in LD (r2 ≥ 0.5) with the top associated variant from the stage 2 meta-analysis for each region ±500kb of the ends of the LD blocks (as expression quantitative trait loci (eQTL) controlled genes are typically less than 500kb from their controlling variant^24^) were input into the UCSC genome browser’s ‘Table Browser’ for RefSeq^25^ and GENCODEv24^26^ genes at each associated locus.

### Human brain gene expression and eQTL analysis

To identify potential functional risk gene(s) at each associated locus we first identified variants with suggestive significance (P>10^−5^) in LD (r2 ≥ 0.5) and within 500kb of the sentinel variants for the 23 associated loci (excluding *APOE*) (N=3,576 variants). We then identified functionally interesting variants in this set of variants using ReguomeDB^22^, HaploReg v4.1^27,28^, GWAS3D^21^. Variants with a RegulomeDB score ≥ 2 (N=160), in high LD (r2 > 0.8) and with evidence of at least one cis-eQTL in any tissue via HaploReg (N=3,407), or with a P ≥ 5 × 10^−8^ in GWAS3D (N=1,120) were selected. We then searched for genes functionally linked via eQTLs in blood (including all immune-related cell types) and brain tissue types using this expanded list of variants (N=3,470). eQTL databases searched included BRAINEAC^29^, SCANdb^30^, the NESDA NTR Conditional eQTL Catalog^31^, GTEx^32^, exSNP^33^ and Zou et al.^34^. Additional eQTL analysis was conducted with INFERNO^35^, where 44 GTEx tissues were searched, with prioritization on the INFERNO tissue classes of brain, blood, and connective tissue (including fibroblasts). INFERNO analyses identified 1,338 unique variants in LD (r2 ≥ 0.7) with the sentinel variants, 1,087 of which are eQTLs (**Supplementary Table 10**).

We also evaluated gene expression of all candidate genes in the associated loci, defined as all genes within ±500kb of the sentinel variant linkage disequilibrium (LD) regions (r^2^ ≥ 0.5) (see **Supplementary Table 7** for a complete list of genes searched), using gene expression data from AlzBase^36^ and the Barres Human and Mouse Brain RNA-Seq Resource^37,38^. AlzBase includes transcription data from brain and blood from aging, non-dementia, mild cognitive impairment, early stage AD and late stage AD. Please see ALZBase (http://alz.big.ac.cn/alzBase/Document) for a complete list of studies included in the search. Genes differentially expressed in the same direction in two or more studies of AD are highlighted in **Supplementary Table 12**.

### Pathway Analysis

Pathway analyses were performed with MAGMA^39^, which performs SNP-wise gene analysis of summary statistics with correction for LD between variants and genes to test whether sets of genes are jointly associated with a phenotype (i.e. LOAD), compared to other genes across the genome Adaptive permutation was used to produce an empirical p-value and an FDR-corrected q-value. Gene-sets used in the analyses were from GO^40,41^, KEGG^42,43^, REACTOME^44,45^, BIOCARTA, and MGI^46^ pathways. Analyses were restricted to gene sets containing between 10 and 500 genes, a total of 10,861 sets. Variants were restricted to common variants (MAF≥0.01) and rare variants (MAF<0.01) only for each analysis, and separate analyses for each model included and excluded the *APOE* region (Chr19:45,116.911-46,318,605). Analyses were also perf12ormed after removal of all genome-wide significant genes. Primary analyses used a 35-kb upstream/10-kb downstream window around each gene in order to potential regulatory variants for each gene, while secondary analyses was run using a 0-kb window^47^. To test for significant correlation between common and rare variant gene results we performed a gene property analysis in MAGMA, regressing the gene-wide association statistics from rare variants on the corresponding statistics from common variants, correcting for LD between variants and genes using the ADGC reference panel. The Aβ-centered network pathway analysis used a curated list of Aβ processing related genes from Campion et al.^48^ Thirty-two Aβ–related gene sets and all 335 genes combined (see Campion et al.^48^ for details) were run in MAGMA pathway analysis on both common (MAF ≥ 0.01) and rare (MAF < 0.01) variant summary results. The combined dataset of 37,635 individuals from the ADGC were used as a reference set for LD calculations in these analyses.

### Validation of prioritization method

Evaluation of the prioritization of the risk genes in genome-wide loci was done using STRINGdb^49^, Jensen Diseases^50^, Jensen Tissues^51^, and the ARCHS4^52^ resource via the EnrichR^53^ tool. We evaluated both the 469 genes set list and the prioritized 68 genes set list (adding in *APOE* to both lists) using the standard settings for both STRINGdb and EnrichR.

## Data Availability

Stage 1 data (individual level) for the GERAD cohort can be accessed by applying directly to Cardiff University. Stage 1 ADGC data are deposited in a NIAGADS- and NIA/NIH-sanctioned qualified-access data repository. Stage 1 CHARGE data are accessible by applying to dbGaP for all US cohorts and to Erasmus University for Rotterdam data. AGES primary data are not available owing to Icelandic laws. Genome-wide summary statistics for the Stage 1 discovery are available from The National Institute on Aging Genetics of Alzheimer’s Disease (NIAGADS) website (https://www.niagads.org/). Stage 2 and stage 3 primary data are available upon request.

## References

1. Thies, W., Bleiler, L. & Alzheimer’s, A. 2013 Alzheimer’s disease facts and figures. Alzheimers Dement 9, 208–245 (2013).

2. Ridge, P. G. et al. Assessment of the genetic variance of late-onset Alzheimer’s disease. Neurobiol. Aging 1–8 (2016). doi:10.1016/j.neurobiolaging.2016.02.024

3. Seshadri, S. et al. Genome-wide analysis of genetic loci associated with Alzheimer disease. JAMA 303, 1832–1840 (2010).

4. Jun, G. et al. Meta-analysis confirms CR1, CLU, and PICALM as alzheimer disease risk loci and reveals interactions with APOE genotypes. Arch Neurol 67, 1473–1484 (2010).

5. Naj, A. C. et al. Common variants at MS4A4/MS4A6E, CD2AP, CD33 and EPHA1 are associated with late-onset Alzheimer’s disease. Nat Genet 43, 436–441 (2011).

6. Harold, D. et al. Genome-wide association study identifies variants at CLU and PICALM associated with Alzheimer’s disease. Nat Genet 41, 1088–1093 (2009).

7. Hollingworth, P. et al. Common variants at ABCA7, MS4A6A/MS4A4E, EPHA1, CD33 and CD2AP are associated with Alzheimer’s disease. Nat Genet 43, 429–435 (2011).

8. Lambert, J. C. et al. Genome-wide association study identifies variants at CLU and CR1 associated with Alzheimer’s disease. Nat Genet 41, 1094–1099 (2009).

9. Lambert, J. C. et al. Meta-analysis of 74,046 individuals identifies 11 new susceptibility loci for Alzheimer’s disease. Nat. Genet. 45, 1452–8 (2013).

10. Gatz, M. et al. Role of genes and environments for explaining Alzheimer disease. Arch Gen Psychiatry 63, 168–174 (2006).

11. Guerreiro, R. et al. TREM2 variants in Alzheimer’s disease. N. Engl. J. Med. 368, 117–27 (2013).

12. Jonsson, T. et al. Variant of TREM2 associated with the risk of Alzheimer’s disease. N. Engl. J. Med. 368, 107–16 (2013).

13. Sims, R. C. et al. Novel rare coding variants in PLCG2, ABI3 and TREM2 implicate microglial-mediated innate immunity in Alzheimer’s disease. Nat. Genet. In press, 1373–1387 (2017).

14. Vassar, R. ADAM10 prodomain mutations cause late-onset Alzheimer’s disease: not just the latest FAD. Neuron 80, 250–253 (2013).

15. Kim, M. et al. Potential late-onset Alzheimer’s disease-associated mutations in the ADAM10 gene attenuate {alpha}-secretase activity. Hum Mol Genet 18, 3987–3996 (2009).

16. Kehoe, P. G. et al. Variation in DCP1, encoding ACE, is associated with susceptibility to Alzheimer disease. Nat. Genet. 21, 71–72 (1999).

17. Meng, Y. et al. Association of polymorphisms in the Angiotensin-converting enzyme gene with Alzheimer disease in an Israeli Arab community. Am J Hum Genet 78, 871–877 (2006).

18. Lehmann, D. J. et al. Large meta-analysis establishes the ACE insertion-deletion polymorphism as a marker of Alzheimer’s disease. Am. J. Epidemiol. 162, 305–17 (2005).

19. Wang, X.-B. et al. Angiotensin-Converting Enzyme Insertion/Deletion Polymorphism Is Not a Major Determining Factor in the Development of Sporadic Alzheimer Disease: Evidence from an Updated Meta-Analysis. PLoS One 9, e111406 (2014).

20. Cai, G. et al. Evidence against a role for rare ADAM10 mutations in sporadic Alzheimer disease. Neurobiol Aging 33, 416–417 e3 (2012).

21. Belbin, O. et al. A multi-center study of ACE and the risk of late-onset Alzheimer’s disease. J Alzheimers Dis 24, 587–597 (2011).

22. Wakutani, Y. et al. Genetic analysis of vascular factors in Alzheimer’s disease. Ann. N. Y. Acad. Sci. 977, 232–8 (2002).

23. Yang, J. et al. Conditional and joint multiple-SNP analysis of GWAS summary statistics identifies additional variants influencing complex traits. Nat. Genet. 44, 369–75, S1-3 (2012).

24. Zhang, Y. et al. An RNA-sequencing transcriptome and splicing database of glia, neurons, and vascular cells of the cerebral cortex. J. Neurosci. 34, 11929–11947 (2014).

25. Zhang, Y. et al. Purification and Characterization of Progenitor and Mature Human Astrocytes Reveals Transcriptional and Functional Differences with Mouse. Neuron 89, 37–53 (2016).

26. Corder, E. H. et al. Protective effect of apolipoprotein E type 2 allele for late onset Alzheimer disease. Nat. Genet. 7, 180–184 (1994).

27. Kim, J., Basak, J. M. & Holtzman, D. M. The role of apolipoprotein E in Alzheimer’s disease. Neuron 63, 287–303 (2009).

28. Steinberg, S. et al. Loss-of-function variants in ABCA7 confer risk of Alzheimer’s disease. Nat. Genet. 26–29 (2015). doi:10.1038/ng.3246

29. Vasquez, J. B., Fardo, D. W. & Estus, S. ABCA7 expression is associated with Alzheimer’s disease polymorphism and disease status. Neurosci Lett 556, 58–62 (2013).

30. Chapuis, J. et al. Increased expression of BIN1 mediates Alzheimer genetic risk by modulating tau pathology. Mol Psychiatry (2013). doi:10.1038/mp.2013.1

31. Rogaeva, E. et al. The neuronal sortilin-related receptor SORL1 is genetically associated with Alzheimer disease. Nat Genet 39, 168–177 (2007).

32. Vardarajan, B. N. et al. Coding mutations in SORL 1 and Alzheimer disease. Ann. Neurol. 77, 215–227 (2015).

33. Suh, J. et al. ADAM10 missense mutations potentiate beta-amyloid accumulation by impairing prodomain chaperone function. Neuron 80, 385–401 (2013).

34. Huang, K. et al. A common haplotype lowers PU.1 expression in myeloid cells and delays onset of Alzheimer&#039;s disease. bioRxiv 20, 1–9 (2017).

35. Brouwers, N. et al. Alzheimer risk associated with a copy number variation in the complement receptor 1 increasing C3b/C4b binding sites. Mol Psychiatry 17, 223–233 (2012).

36. Flister, M. J. et al. Identifying multiple causative genes at a single GWAS locus. Genome Res 23, 1996–2002 (2013).

37. Bai, Z. et al. AlzBase: an Integrative Database for Gene Dysregulation in Alzheimer’s Disease. Mol. Neurobiol. 53, 310–319 (2016).

38. de Leeuw, C. A., Mooij, J. M., Heskes, T. & Posthuma, D. MAGMA: Generalized Gene-Set Analysis of GWAS Data. PLoS Comput. Biol. 11, 1–19 (2015).

39. Stefanis, L. alpha-Synuclein in Parkinson’s disease. Cold Spring Harb. Perspect. Med. 2, 1–23 (2012).

40. Takeda, A. et al. C-terminal alpha-synuclein immunoreactivity in structures other than Lewy bodies in neurodegenerative disorders. Acta Neuropathol. 99, 296–304 (2000).

41. Campion, D., Pottier, C., Nicolas, G., Le Guennec, K. & Rovelet-Lecrux, A. Alzheimer disease: modeling an Aβ-centered biological network. Mol. Psychiatry 1–11 (2016). doi:10.1038/mp.2016.38

42. Gandy, S. & Heppner, F. L. Microglia as dynamic and essential components of the amyloid hypothesis. Neuron 78, 575–577 (2013).

43. Sims, R., van der Lee, S. J., Naj, A. C., Bellenguez, C. & et al. Novel rare coding variants in PLCG2, ABI3 and TREM2 implicate microglial-mediated innate immunity in Alzheimer’s disease. Nat. Genet 49, 1373–1383 (2017).

44. Haass, C., Kaether, C., Thinakaran, G. & Sisodia, S. Trafficking and proteolytic processing of APP. Cold Spring Harb Perspect Med 2, a006270 (2012).

45. Kleinberger, G. et al. TREM2 mutations implicated in neurodegeneration impair cell surface transport and phagocytosis. Sci. Transl. Med. 6, (2014).

46. Postina, R. et al. A disintegrin-metalloproteinase prevents amyloid plaque formation and hippocampal defects in an Alzheimer disease mouse model. J. Clin. Invest. 113, 1456–1464 (2004).

47. Kurabayashi, N., Nguyen, M. D. & Sanada, K. The G protein-coupled receptor GPRC5B contributes to neurogenesis in the developing mouse neocortex. Development 140, 4335–4346 (2013).

48. Cool, B. H. et al. A flanking gene problem leads to the discovery of a Gprc5b splice variant predominantly expressed in C57BL/6J mouse brain and in maturing neurons. PLoS One 5, (2010).

49. Kim, Y.-J., Sano, T., Nabetani, T., Asano, Y. & Hirabayashi, Y. GPRC5B Activates Obesity-Associated Inflammatory Signaling in Adipocytes. Sci. Signal. 5, ra85–ra85 (2012).

50. Bhat, K. et al. The 19S proteasome ATPase Sug1 plays a critical role in regulating MHC class II transcription. Mol. Immunol. 45, 2214–24 (2008).

51. Inostroza-Nieves, Y., Venkatraman, P. & Zavala-Ruiz, Z. Role of Sug1, a 19S proteasome ATPase, in the transcription of MHC I and the atypical MHC II molecules, HLA-DM and HLA-DO. Immunol. Lett. 147, 67–74 (2012).

52. Farrer, L. A. et al. Association Between Angiotensin-Converting Enzyme and Alzheimer Disease. New Engl. J. Med. 57, 210–14 (2000).

53. Miners, J. S. et al. Angiotensin-converting enzyme levels and activity in Alzheimer’s disease: Differences in brain and CSF ACE and association with ACE1 genotypes. Am. J. Transl. Res. 1, 163–177 (2009).

54. Jochemsen, H. M. et al. The association of angiotensin-converting enzyme with biomarkers for Alzheimer’s disease. Alzheimer’s Res. Ther. 6, 1–10 (2014).

55. Kauwe, J. S. K. et al. Genome-Wide Association Study of CSF Levels of 59 Alzheimer’s Disease Candidate Proteins: Significant Associations with Proteins Involved in Amyloid Processing and Inflammation. PLoS Genet. 10, e1004758 (2014).

56. Baranello, R. J. et al. Amyloid-beta protein clearance and degradation (ABCD) pathways and their role in Alzheimer’s disease. Curr. Alzheimer Res. 12, 32–46 (2015).

57. Kehoe, P. G. The Coming of Age of the Angiotensin Hypothesis in Alzheimer’s Disease: Progress Toward Disease Prevention and Treatment? J. Alzheimer’s Dis. 62, In Press (2018).

58. Kehoe, P. G. et al. The Rationale and Design of the Reducing Pathology in Alzheimer’s Disease through Angiotensin TaRgeting (RADAR) Trial. J. Alzheimer’s Dis. 61, 803–814 (2017).

59. Miguel, R. F., Pollak, A. & Lubec, G. Metalloproteinase ADAMTS-1 but not ADAMTS-5 is manifold overexpressed in neurodegenerative disorders as Down syndrome, Alzheimer’s and Pick’s disease. Brain Res. Mol. Brain Res. 133, 1–5 (2005).

60. Suttkus, A. et al. Aggrecan, link protein and tenascin-R are essential components of the perineuronal net to protect neurons against iron-induced oxidative stress. Cell Death Dis. 5, e1119 (2014).

61. Végh, M. J. et al. Reducing hippocampal extracellular matrix reverses early memory deficits in a mouse model of Alzheimer’s disease. Acta Neuropathol. Commun. 2, 76 (2014).

62. Morawski, M., Filippov, M., Tzinia, A., Tsilibary, E. & Vargova, L. ECM in brain aging and dementia. Prog. Brain Res. 214, 207–27 (2014).

63. Wilcock, D. M. Neuroinflammation in the aging down syndrome brain; lessons from Alzheimer’s disease. Curr Gerontol Geriatr Res 2012, 170276 (2012).

64. Dourlen, P. et al. Functional screening of Alzheimer risk loci identifies PTK2B as an in vivo modulator and early marker of Tau pathology. Mol. Psychiatry 22, 874–883 (2017).

65. Chapuis, J. et al. Genome-wide, high-content siRNA screening identifies the Alzheimer’s genetic risk factor FERMT2 as a major modulator of APP metabolism. Acta Neuropathol. 133, 955–966 (2017).

66. Shulman, J. M. et al. Functional screening in Drosophila identifies Alzheimer’s disease susceptibility genes and implicates tau-mediated mechanisms. Hum. Mol. Genet. 23, 870–877 (2014).

67. Murray, M. E. et al. Clinicopathologic and 11C-Pittsburgh compound B implications of Thal amyloid phase across the Alzheimer’s disease spectrum. Brain 1–12 (2015). doi:10.1093/brain/awv050

68. Shi, Y. et al. ApoE4 markedly exacerbates tau-mediated neurodegeneration in a mouse model of tauopathy. Nature 549, 523–527 (2017).

69. Brier, M. R. et al. Tau and Aβ imaging, CSF measures, and cognition in Alzheimer’s disease. Sci. Transl. Med. 8, 338ra66 (2016).

## Methods References

1. Genomes Project, C. et al. An integrated map of genetic variation from 1,092 human genomes. Nature 491, 56–65 (2012).

2. Howie, B. N., Donnelly, P. & Marchini, J. A flexible and accurate genotype imputation method for the next generation of genome-wide association studies. PLoS Genet 5, e1000529 (2009).

3. Delaneau, O., Marchini, J. & Zagury, J. F. A linear complexity phasing method for thousands of genomes. Nat Methods 9, 179–181 (2012).

4. Li, Y., Willer, C. J., Ding, J., Scheet, P. & Abecasis, G. R. MaCH: using sequence and genotype data to estimate haplotypes and unobserved genotypes. Genet Epidemiol 34, 816–834 (2010).

5. Howie, B., Fuchsberger, C., Stephens, M., Marchini, J. & Abecasis, G. R. Fast and accurate genotype imputation in genome-wide association studies through pre-phasing. Nat Genet 44, 955–959 (2012).

6. Howie, B., Marchini, J. & Stephens, M. Genotype imputation with thousands of genomes. G3 1, 457–470 (2011).

7. Lambert, J. C. et al. Meta-analysis of 74,046 individuals identifies 11 new susceptibility loci for Alzheimer’s disease. Nat. Genet. 45, 1452–8 (2013).

8. Price, A. L. et al. Principal components analysis corrects for stratification in genome-wide association studies. Nat Genet 38, 904–909 (2006).

9. Ma, C. et al. Recommended Joint and Meta-Analysis Strategies for Case-Control Association Testing of Single Low-Count Variants. Genet Epidemiol 37, 539–550 (2013).

10. Chen, M.-H. H. & Yang, Q. GWAF: an R package for genome-wide association analyses with family data. Bioinformatics 26, 580–581 (2010).

11. Willer, C. J., Li, Y. & Abecasis, G. R. METAL: fast and efficient meta-analysis of genomewide association scans. Bioinformatics 26, 2190–2191 (2010).

12. Lee, S., Abecasis, G. R., Boehnke, M. & Lin, X. Rare-Variant Association Analysis: Study Designs and Statistical Tests. Am. J. Hum. Genet. 95, 5–23 (2014).

13. Pruim, R. J. et al. LocusZoom: regional visualization of genome-wide association scan results. Bioinformatics 26, 2336–2337 (2010).

14. Yang, J. et al. Conditional and joint multiple-SNP analysis of GWAS summary statistics identifies additional variants influencing complex traits. Nat Genet 44, 369–75, S1-3 (2012).

15. Machiela, M. J. & Chanock, S. J. LDlink : A web-based application for exploring population-specific haplotype structure and linking correlated alleles of possible functional variants. 1–3 (2015).

16. Patterson, N., Price, A. L. & Reich, D. Population structure and eigenanalysis. PLoS Genet 2, e190 (2006).

17. Ritchie, G. R., Dunham, I., Zeggini, E. & Flicek, P. Functional annotation of noncoding sequence variants. Nat Methods 11, 294–296 (2014).

18. Kircher, M. et al. A general framework for estimating the relative pathogenicity of human genetic variants. Nat Genet 46, 310–315 (2014).

19. Wang, K., Li, M. & Hakonarson, H. ANNOVAR: functional annotation of genetic variants from high-throughput sequencing data. Nucleic Acids Res 38, e164 (2010).

20. McLaren, W. et al. Deriving the consequences of genomic variants with the Ensembl API and SNP Effect Predictor. Bioinformatics 26, 2069–2070 (2010).

21. Li, M. J., Wang, L. Y., Xia, Z., Sham, P. C. & Wang, J. GWAS3D: detecting human regulatory variants by integrative analysis of genome-wide associations, chromosome interactions and histone modifications. Nucleic Acids Res 41, W150–W158 (2013).

22. Boyle, A. P. et al. Annotation of functional variation in personal genomes using RegulomeDB. Genome Res 22, 1790–1797 (2012).

23. Andersson, R. et al. An atlas of active enhancers across human cell types and tissues. Nature 507, 455–461 (2014).

24. Zhang, X. et al. Synthesis of 53 tissue and cell line expression QTL datasets reveals master eQTLs. BMC Genomics 15, 532 (2014).

25. Pruitt, K. D., Tatusova, T., Brown, G. R. & Maglott, D. R. NCBI Reference Sequences (RefSeq): current status, new features and genome annotation policy. Nucleic Acids Res 40, D130–5 (2012).

26. Harrow, J. et al. GENCODE: the reference human genome annotation for The ENCODE Project. Genome Res 22, 1760–1774 (2012).

27. Ward, L. D. & Kellis, M. HaploReg: a resource for exploring chromatin states, conservation, and regulatory motif alterations within sets of genetically linked variants. Nucleic Acids Res 40, D930–4 (2012).

28. Ward, L. D. & Kellis, M. HaploReg v4: systematic mining of putative causal variants, cell types, regulators and target genes for human complex traits and disease. Nucleic Acids Res. 44, gkv1340 (2015).

29. Ramasamy, A. et al. Genetic variability in the regulation of gene expression in ten regions of the human brain. Nat. Neurosci. 17, 1418–1428 (2014).

30. Gamazon, E. R. et al. SCAN: SNP and copy number annotation. Bioinformatics 26, 259–262 (2010).

31. Jansen, R. et al. Conditional eQTL Analysis Reveals Allelic Heterogeneity of Gene Expression. Hum. Mol. Genet. 26, 1444–1451 (2017).

32. Consortium, G. Te. The Genotype-Tissue Expression (GTEx) project. Nat Genet 45, 580–585 (2013).

33. Yu, C.-H., Pal, L. R. & Moult, J. Consensus Genome-Wide Expression Quantitative Trait Loci and Their Relationship with Human Complex Trait Disease. Omi. A J. Integr. Biol. 20, 400–414 (2016).

34. Zou, F. et al. Brain expression genome-wide association study (eGWAS) identifies human disease-associated variants. PLoS Genet 8, e1002707 (2012).

35. Amlie-Wolf, A. et al. INFERNO – INFERring the molecular mechanisms of NOncoding genetic variants. bioRxiv Oct 30, (2017).

36. Bai, Z. et al. AlzBase: an Integrative Database for Gene Dysregulation in Alzheimer’s Disease. Mol. Neurobiol. 53, 310–319 (2016).

37. Zhang, Y. et al. An RNA-sequencing transcriptome and splicing database of glia, neurons, and vascular cells of the cerebral cortex. J. Neurosci. 34, 11929–11947 (2014).

38. Zhang, Y. et al. Purification and Characterization of Progenitor and Mature Human Astrocytes Reveals Transcriptional and Functional Differences with Mouse. Neuron 89, 37–53 (2016).

39. de Leeuw, C. A., Mooij, J. M., Heskes, T. & Posthuma, D. MAGMA: Generalized Gene-Set Analysis of GWAS Data. PLoS Comput. Biol. 11, 1–19 (2015).

40. Ashburner, M. et al. Gene ontology: tool for the unification of biology. The Gene Ontology Consortium. Nat Genet 25, 25–29 (2000).

41. Blake, J. A. et al. Gene ontology consortium: Going forward. Nucleic Acids Res. 43, D1049–D1056 (2015).

42. Kanehisa, M., Sato, Y., Kawashima, M., Furumichi, M. & Tanabe, M. KEGG as a reference resource for gene and protein annotation. Nucleic Acids Res. 44, D457–D462 (2016).

43. Ogata, H. et al. KEGG: Kyoto encyclopedia of genes and genomes. Nucleic Acids Res. 27, 29–34 (1999).

44. Fabregat, A. et al. The reactome pathway knowledgebase. Nucleic Acids Res. 44, D481–D487 (2016).

45. Croft, D. et al. Reactome: a database of reactions, pathways and biological processes. Nucleic Acids Res 39, D691–7 (2011).

46. Eppig, J. T., Blake, J. a, Bult, C. J., Kadin, J. a & Richardson, J. E. The Mouse Genome Database (MGD): facilitating mouse as a model for human biology and disease. Nucleic Acids Res. 1–11 (2014). doi:10.1093/nar/gku967

47. Network, P. Psychiatric genome-wide association study analyses implicate neuronal, immune and histone pathways. Nat. Neurosci. (2015). doi:10.1038/nn.3922

48. Campion, D., Pottier, C., Nicolas, G., Le Guennec, K. & Rovelet-Lecrux, A. Alzheimer disease: modeling an Aβ-centered biological network. Mol. Psychiatry 1–11 (2016). doi:10.1038/mp.2016.38

49. Szklarczyk, D. et al. STRING v10: Protein-protein interaction networks, integrated over the tree of life. Nucleic Acids Res. 43, D447–D452 (2015).

50. Pletscher-Frankild, S., Pallejà, A., Tsafou, K., Binder, J. X. & Jensen, L. J. DISEASES: Text mining and data integration of disease-gene associations. Methods 74, 83–89 (2015).

51. Santos, A. et al. Comprehensive comparison of large-scale tissue expression datasets. PeerJ 3, e1054 (2015).

52. Lachmann, A. et al. Massive Mining of Publicly Available RNA-seq Data from Human and Mouse. Bioarxiv 1–9 (2017). doi:10.1101/189092

53. Kuleshov, M. V. et al. Enrichr: a comprehensive gene set enrichment analysis web server 2016 update. Nucleic Acids Res. 44, W90–W97 (2016).

